# Phenomics data processing: A plot-level model for repeated measurements to extract the timing of key stages and quantities at defined time points

**DOI:** 10.1101/2021.05.02.442243

**Authors:** Lukas Roth, María Xosé Rodríguez-Álvarez, Fred van Eeuwijk, Hans-Peter Piepho, Andreas Hund

**Affiliations:** ETH Zurich, Institute of Agricultural Sciences, Universitätstrasse 2, 8092 Zurich, Switzerland; BCAM – Basque Center for Applied Mathematics, Alameda de Mazarredo, 14. E-48009 Bilbao, Basque Country, Spain; IKERBASQUE, Basque Foundation for Science, Bilbao, Spain; Wageningen University and Research, Biometris, P.O. Box 16, 6700 AA Wageningen, The Netherlands; University of Hohenheim, Institute for Crop Science, Biostatistics Unit, Fruwirthstrasse 23, 70593 Stuttgart, Germany

**Keywords:** High-throughput field phenotyping, dynamic modeling, stage-wise processing, canopy height

## Abstract

Decision-making in breeding increasingly depends on the ability to capture and predict crop responses to changing environmental factors. Advances in crop modeling as well as high-throughput field phenotyping (HTFP) hold promise to provide such insights. Processing HTFP data is an interdisciplinary task that requires broad knowledge on experimental design, measurement techniques, feature extraction, dynamic trait modeling, and prediction of genotypic values using statistical models. To get an overview of sources of variation in HTFP, we develop a general plot-level model for repeated measurements. Based on this model, we propose a seamless step-wise procedure that allows for carry on of estimated means and variances from stage to stage. The process builds on the extraction of three intermediate trait categories; (1) timing of key stages, (2) quantities at defined time points or periods, and (3) dose-response curves. In a first stage, these intermediate traits are extracted from low-level traits’ time series (e.g., canopy height) using P-splines and the quarter of maximum elongation rate method (QMER), as well as final height percentiles. In a second and third stage, extracted traits are further processed using a stage-wise linear mixed model analysis. Using a wheat canopy growth simulation to generate canopy height time series, we demonstrate the suitability of the stage-wise process for traits of the first two above-mentioned categories. Results indicate that, for the first stage, the P-spline/QMER method was more robust than the percentile method. In the subsequent two-stage linear mixed model processing, weighting the second and third stage with error variance estimates from the previous stages improved the root mean squared error. We conclude that processing phenomics data in stages represents a feasible approach if estimated means and variances are carried forward from one processing stage to the next. P-splines in combination with the QMER method are suitable tools to extract timing of key stages and quantities at defined time points from HTFP data.

**Highlights:**

- General plot-level model for repeated high-throughput field phenotyping measurements
- Extraction of three main intermediate trait categories for dynamic modeling
- Seamless processing approach that integrates temporal and spatial modeling
- Phenomics data processing cheatsheet

## 1. Introduction

Advances in high-throughput field phenotyping (HTFP) allow for capture of large data sets with high temporal and spatial resolution (Rebetzke et al., 2019). Summarizing these spatio-temporal data in a meaningful way is essential to support selection and decision-making in breeding. In HTFP the primary data often consists of images, point measurements, orthophotos, or point clouds from which low-level traits (e.g., shoot counts, canopy cover, canopy height, or senescence) are extracted. After feature extraction, these low-level traits may be tracked over time in a subsequent temporal modelling step (van Eeuwijk et al., 2019;Moreira et al., 2020). If monitored across the lifetime of a plant, low-level traits often follow some sort of monotonically increasing function (e.g., canopy height or senescence) or concave function (e.g., number of growing shoots or canopy cover), which allows for estimates of dynamics’ characteristics. These estimates are referred to as intermediate traits.

Estimating such intermediate traits from spatio-temporal measurements implies *a priori* knowledge of growth processes, best summarized in crop growth models. Crop models have rapidly gained in complexity over time, culminating in the description of plants by 3-D functional-structural models (Vos et al., 2010). Indoor platforms have proven useful to provide genotype-specific parameter estimates for such models (Tardieu et al., 2017), but discrepancies between field and indoor experiments raised doubts if results are always directly transferable (Poorter et al., 2016). Field-based phenotyping may allow assessing and improving the performance of such crop models (Ramirez-Villegas et al., 2015), but also provide genotype-specific parameter estimates that are better transferable to real-world conditions (Araus et al., 2018).

While under controlled conditions environmental factors may be adequately controlled, the lack of control over meteorological conditions poses a major challenge for field phenotyping. Several additional types of errors need to be considered, which can be classified into those directly affecting the sensor reading, and those affecting the plant development.

In HTFP there are attempts to quantify genotype-specific timing of phenology stages (Hurtado et al., 2012) and response patterns to distinct environmental variables like temperature (Grieder et al., 2015;Kronenberg et al.,2020a). A comparable approach in genomics uses functional mapping of quantitative trait loci (QTLs), e.g., based on logistic growth curves (Ma et al., 2002;Malosetti et al., 2006). Ma et al. proposed to distinguish three biological processes in such models: allometric laws, growth models, and reaction norms. Characterizing growth dynamics using field data becomes increasingly difficult as models become more complex. A solution is to predict crop growth from arbitrary traits or scores that lack a clear physiological interpretation. In phenomics, this was demonstrated using serial measurements as predictors for statistical learning (Ubbens et al., 2020;Maimaitijiang et al., 2020;Herrero-Huerta et al., 2020). In genomics, comparable approaches are based on functional principal component analysis, where curves are specified as linear combinations of basis functions, and the corresponding scores then used as intermediate traits (Kwak et al., 2016;Moreira et al., 2020).

From a plant physiology point of view, such approaches represent a ’black box‘, as drawing conclusions about the biological importance of the underlying traits is difficult. In addition, if the traits or scores that lack a clear physiological interpretation do not correspond to features under genetic control, the resulting statistical models will not account for a sufficient amount of the phenotype variance to be useful in a breeding context. Therefore, we believe that a classical approach to extract traits related to distinct crop ideotypes based on current existing physiological knowledge about the biological basis of the dynamic growth process is more suitable (see also van Eeuwijk et al., 2019;Bustos-Korts et al., 2019), as it enables to connect HTFP observations to expert knowledge in crop physiology acquired over decades by a large scientific community (Hund et al., 2019). This approach may then represent a standard to compare modern learning approaches with.

Based on HTFP literature and the biological processes described in Ma et al. (2002), we identified three main intermediate trait categories which can be related to ideotype concepts:

1. **Timing of key stages**: Turning points in the dynamics of numeric measurements which may be related to phenology; e.g., beginning of stem elongation (Kronenberg et al., 2017), time of canopy closure (Soltani and Galeshi, 2002), time of maximum canopy growth rate (Borra-Serrano et al., 2020), heading and flowering (Sadeghi-Tehran et al., 2017), or onset and end of senescence (Anderegg et al., 2020;Aasen et al.,2020). Genotype-specific responses to environmental covariates and/or indices may help to predict key stages; e.g., flowering time (Millet et al., 2019).
2. **Quantities at defined time points or periods**: Traits based on numeric measurements; either at a steady state; e.g., canopy temperature between flowering and beginning of senescence (Perich et al., 2020), or at well-defined time points; e.g., number of tillers at beginning of stem elongation (Roth et al., 2020) and at harvest (Jin et al., 2019), number of ears at harvest (Fernandez-Gallego et al., 2018), or canopy cover at maximum (Borra-Serrano et al., 2020). Area-under-the-curve traits may represent a special case of this category where one summarizes quantities over a defined range of time points (Blancon et al., 2019).
3. **Dose-response curves**: Traits that describe developmental responses in dependence on covariates between clearly defined boundary key stages, i.e., parameters of curves. Dose-response experiments are classically conducted under controlled conditions, e.g., by examining the response of leaves to temperature and water deficit (Reymond et al., 2003) and to soil water deficiency and evaporative demand (Welcker et al., 2011) during their linear growth phase.Reymond et al. (2003) partially included field based measurements, but more recently, such experiments were conducted completely in the field; e.g., in the early, exponential development phase of canopy cover between emergence and tillering (Grieder et al., 2015) or at the linear development phase of canopy height between start and end of stem elongation (Kronenberg et al., 2020a).

Despite the differences in subsequent processing, the extraction of each of the three different trait categories is a highly repetitive task which requires analysis routines with sufficient robustness and generality. While timing of key stages and quantities belong to growth model processes, dose-response curves relate to reaction norm processes (Via et al., 1995). Arguably, dose-response curves represent the most challenging modelling aspect in field phenotying, as they require quantifying growth and relating it to environmental covariates. We will cover this aspect in a follow-up paper (Roth et al., 2021). However, a robust evaluation of such dose-response curves requires determining the boundaries between which a steady development takes place. Here, we aim to develop a method to extract such timing of key stages and quantity traits.

We start by developing a plot-level model for repeated measurements, with a focus on the outdoor field phenotyping platform (FIP) (Kirchgessner et al., 2017). The FIP allows us to densely monitor a large set of replicated genotypes (i.e., two replicates of 350 genotypes) over a whole growing season with genotypes being the only treatment. The aim of such experiments is to i) allow developing new traits and phenotyping methodologies; ii) characterize a specific target environment including the targeted optimal genotype (a so called ideotype, for an overview of definitions see https://kp.ethz.ch/research/research-and-thesis-projects/phys-breeding/glossar.html); and iii) to serve as part of multi-environment trials (MET) that cover a target population of environments, defined as “the ensemble of conditions (including impact from management) that a commercially cultivated crop is likely to experience in a given geographic area” (Chenu et al., 2017).

Processing MET data is often done using linear mixed models (Piepho et al., 2012). Single-stage models that account for within-environment effects and between-environment effects simultaneously are considered the gold standard (Welham et al., 2010). Nevertheless, stage-wise approaches where individual environments are analyzed separately in a first stage are more common because of their simplicity and computational efficiency (Möhring and Piepho, 2009). When using weights in the second stage based on variance estimations for the first stage, such approaches can adequately approximate a single-stage analysis (Piepho et al., 2012).

Here, we propose a possible solution to analyze HTFP experiments based on existing statistical tools such as P-splines (Eilers and Marx, 1996) and stage-wise linear mixed model analysis. We further evaluate and demonstrate the suitability of the approach to extract the timing of key stages and quantities at defined time points from low-level traits using simulated wheat canopy height data.

## 2. Materials and Methods

### 2.1. A plot-level model for repeated measurements

A planned experiment must include an experimental design (Figure 1a+b, green boxes) in which the treatment factors to be tested are randomly assigned to experimental units (usually plots). For simplicity, we will refer to experimental units as plots throughout the manuscript. For the FIP, the only treatment factor are genotypes. Reasoned by the immobility of the FIP, the design comprises only one site but multiple years. The data for each year holds a subset of treatment levels (genotypes) together with checks and design factors (blocks) to allow correcting for spatial variability at the site. In the specific case of the FIP a panel of on average 345 genotypes is replicated twice per year and each replication is planted on a different lot (one of six spatially continuous areas integrated in a crop roation in the FIP area). Each replication is augmented with spatial checks in a 3×3 block arrangement to allow accounting for spatially correlated nuisance factors (see below).

**Figure 1:**
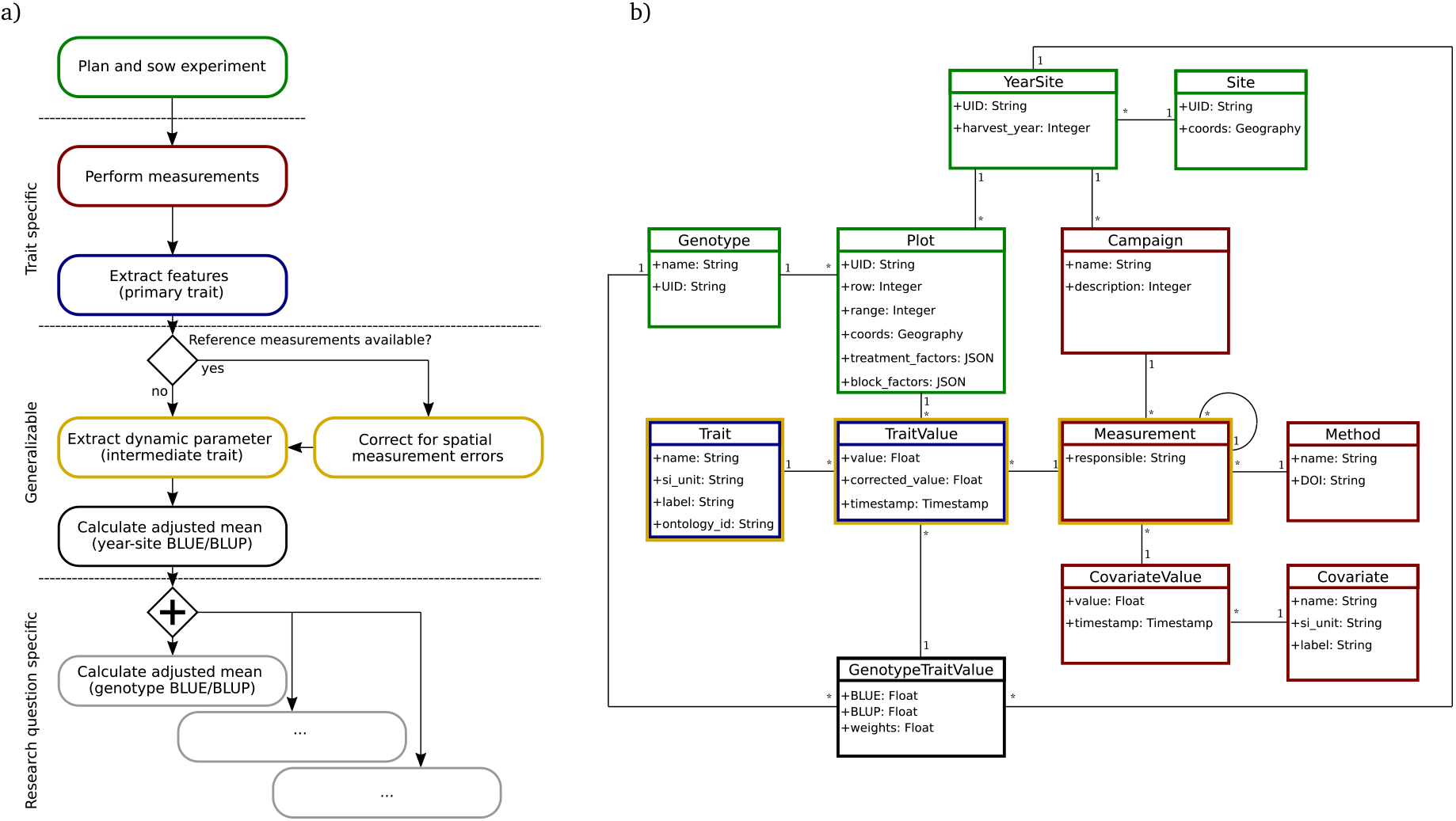
Minimal process-driven model for the FIP: a) Process model, b) Data model.

Performing measurements implies the application of a sensing device collecting measurements from a plot. This process results in data which either directly represent a trait value (e.g., a point measurement of temperature) or can be translated to one or several low-level traits by means of feature-extraction (Figure 1a+b, blue boxes).

A campaign is, for a particular quantity, the repeated collection of its measurements from the same plots over a total interval that might stretch from days to the entire growing season (Figure 1a+b, red boxes). A campaign time point is the interval lasting from seconds to hours (depending on the measurement technology and design size) over which one collection of all plots of an experimental design in a campaign is performed. In contrast, a timestamp is the exact time point at which a quantity is measured in a specific plot.

The same approach of campaigns and measurements also applies to covariate measurements (Figure 1a+b, red boxes), but the level at which the covariate is measured adds additional complexity. The FIP site includes a weather station that logs a standard suite of meteorological variables at 10 minutes intervals. Because the FIP studies annual crops and only has one such station, the covariates it measures reside at the year-site level. On the other extreme, if measuring for example meristem temperatures with thermocouples, the covariates the devices measure reside at the organ level.

The measurement level has consequences on what one regards as phenotypic heterogeneity caused by covariate variation: For the example of the FIP site where the covariates reside at the year-site level, one must consider heterogeneity caused by covariate variations at plot, plant and organ levels and their effects on plant growth (Figure 2a). Namely, these effects include variations of the timing of key stages (Figure 2a1+c1) resulting in growth period condition variations (Figure 2b2+c2) and consequently variations of quantities at defined time points (Figure 2a3+c3).

**Figure 2:**
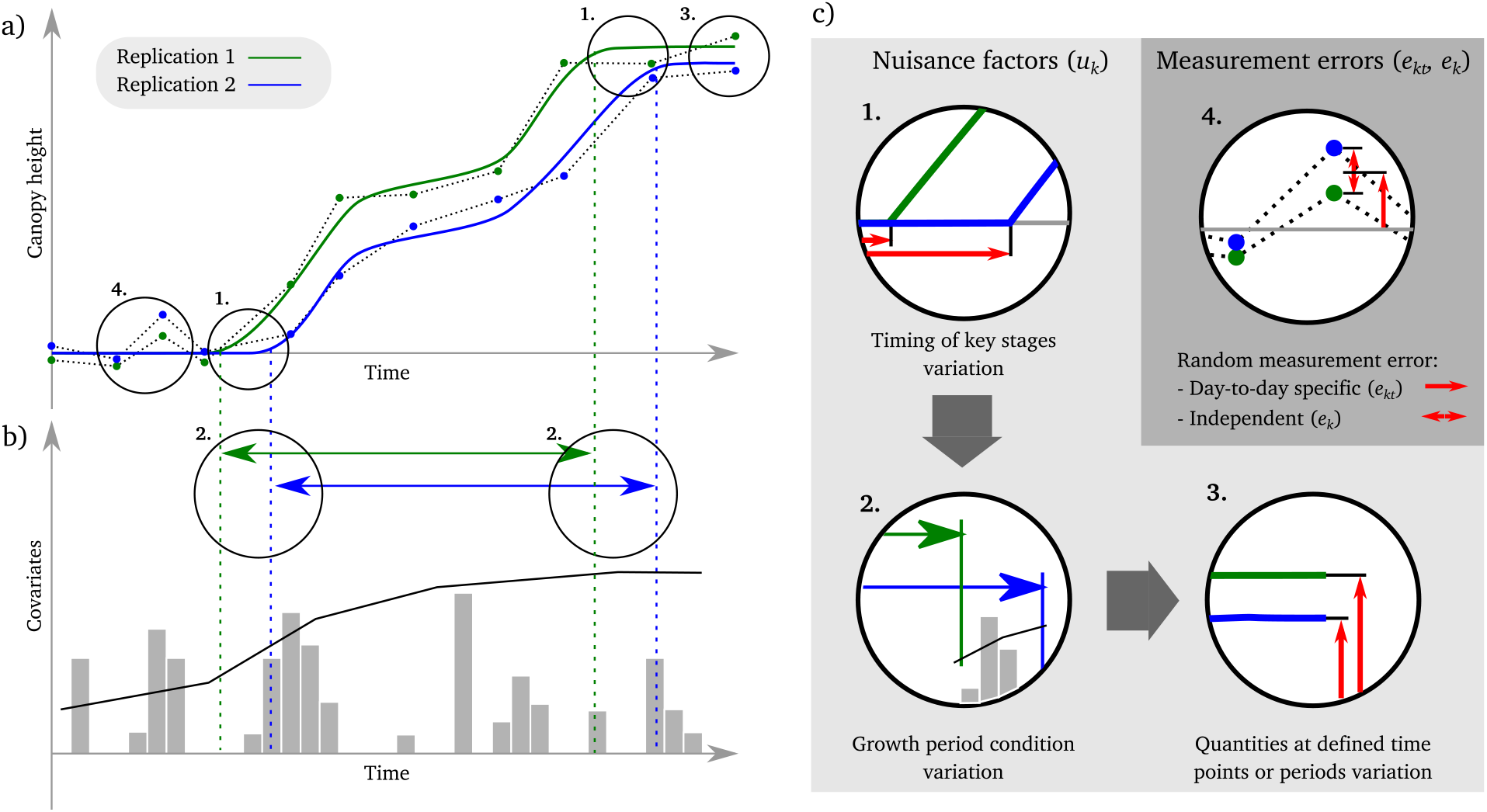
Sources of variation in HTFP on the example of canopy height measurements. (a) Canopy height development of two replications of the same genotype (green and blue lines) and realized measurement time points (green and blue dots). (b) Covariate measurements during the growth phase of canopies (e.g., temperature and precipitation). (c) Sources of variation, the item number 1–4 correspond to the respective numbered items in (a) and (b): (1) spatial and crop-husbandry effect leading to different timings of key stages, e.g., start and end of stem elongation; (2) timing of key stage variations leading to variations in the different gradients of environmental covariates, e.g., temperature gradients in the stem elongation phase; (3) spatial and crop-husbandry effects leading to quantitative variations in trait values; e.g., final height at the end of the stem elongation phase; (4) Day-to-day random measuring errors, e.g., related to differing conditions between measurement days; and independent random measuring errors, e.g., related to the measurement precision of the device.

In a phenotyping experiment one has to distinguish between nuisance factors affecting the growth and development of the plant (Figure 2a1–3), and measurement errors affecting the precision at which a certain phenotype is measured at a given time (Figure 2a4). The latter factors may affect whole campaign time points (i.e., at the day-to-day level, Figure 2c4, red one-sided arrow) but also individual measuring time points within a day (Figure 2c4, red two-sided arrow). Nuisance factors affecting growth and development are, e.g., soil fertility inhomogeneities, spatial temperature gradients, mice, herbivore damages, and other abiotic and biotic factors varying spatially and temporally in the field. By using randomization and blocking in the experimental design, such factors can be accounted for, as was done in this simulation study.

**Figure 3:**
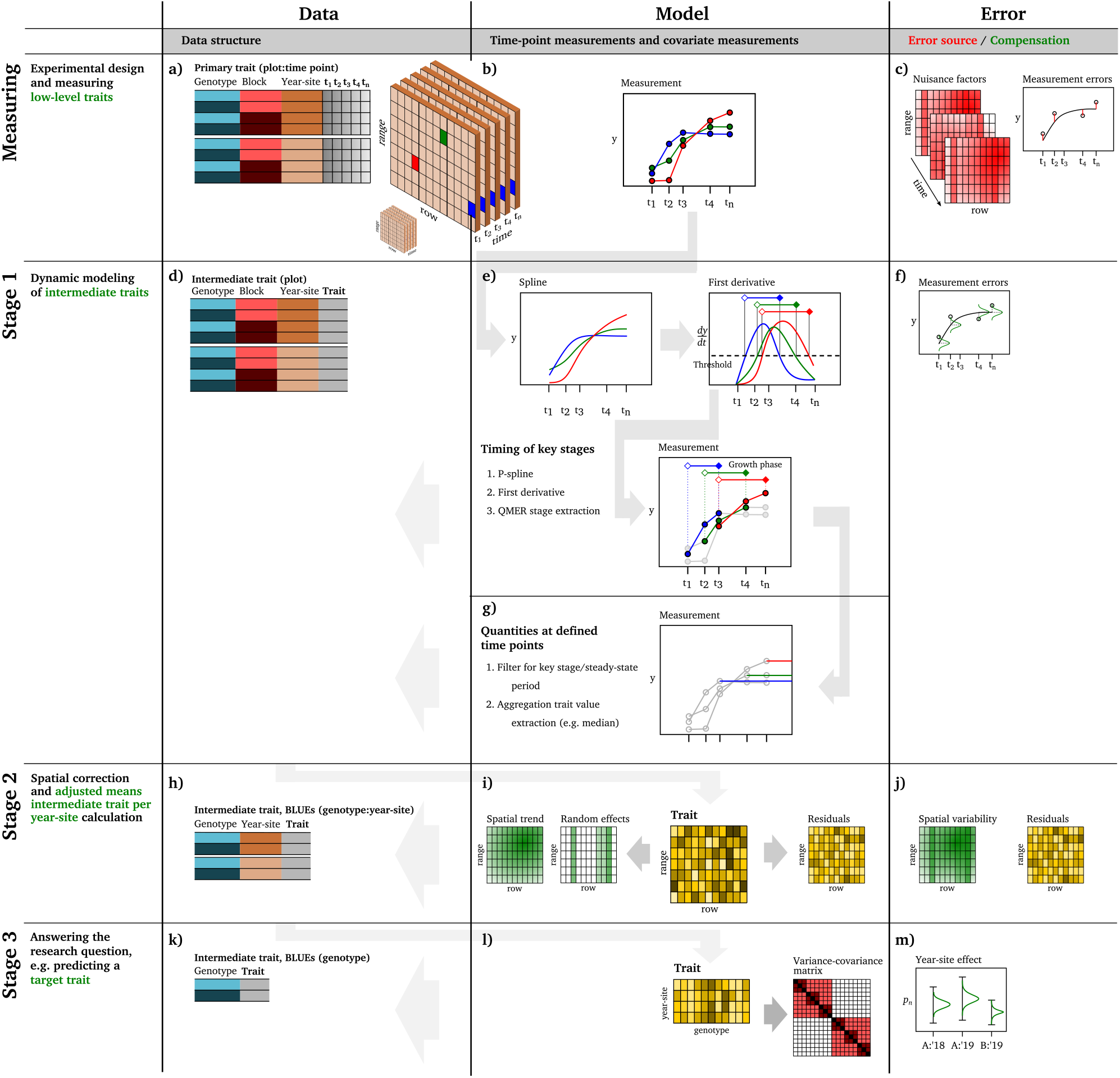
Phenomics data processing cheatsheet: Extraction of timing of key stages and quantities at defined time points from high-throughput field phenotpying measurements (a-c) in three stages. The first stage models the dynamic of plot-based time series to extract plot-based intermediate traits (d-g), the second stage models the spatial context to extract adjusted genotype-based means per year-site (h-j), and the third stage models year-site effects to, e.g., predict a target trait (k-m).

Sources of measurement errors include factors differing between campaign time points. These factors may lead to day-specific under- or overestimation of measurements, arising for example from positioning shifts of the sensor, re-adjustment of sensor settings between measurement campaigns, changes in canopy characteristics after rain or during hot days, or differing illumination conditions (Figure 2c4, red one-sided arrow). One mitigation strategy, when feasible, is to use calibration targets. Apart from the effects related to the whole campaign time point, changing conditions during the measuring sequence may lead to additional, temporally correlated measurement errors among measuring time points. Typical unavoidable sources for such errors are changing weather conditions during the sequential measurement of the first to the last plot allocated to a field design. Thus, usually there is a temporal gradient in the direction of increasing plot number. When such temporal effects are confounded with nuisance factors, the analogous types of strategies mentioned earlier (e.g., blocking, randomizing, and calibration targets) can be applied. Finally, random measurement device errors (Figure 2c4, red two-sided arrow) represent another source of variation in HTFP These errors are usually assumed to be identically and independently normally distributed (i.i.d.).

Consequently, we define a HTFP observation *y_kt_* for the *t*-th time point on the *k*-th plot (*k* = 1,…, *K*) as the result of a dynamic data generating model *g* that is a function of time *t* and of a vector 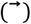 of plot-specific crop growth parameters 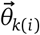 associated with genotype *i* modulated (;) by a vector of time-varying covariates 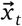, and of a plot residual *e_kt_* that is i.i.d. with a variance that is constant over time 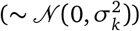,

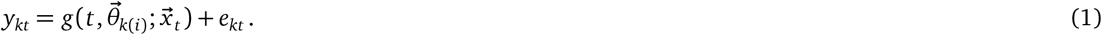

(Figure 3a-c). While *e_kt_* will account for random measurement device errors (“white noise” over space and time), we assume here that *g* will absorb any spatio-temporal correlation among measurements. Dynamic modeling (Equation 1) is done separately for each individual plot-based time series (Stage 1), i.e., (*y_k_*_1_,…, *y_kT_*) (Figure 3d–g), where *T* denotes the last measurement.

Stage 1 therefore allows estimating plot-specific crop growth model parameters 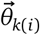. Those crop growth model parameters will become a phenotypic trait when measured / estimated for a set of genotypes. Correcting for spatial correlations is done in a subsequent stage (Stage 2) of this stage-wise approach to obtain estimates of genotype specific crop growth model parameters 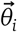 (Figure 3h–j). This estimation step is done separately for each crop growth model parameter *θ_n,i_* in 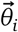 based on fitting the linear mixed model

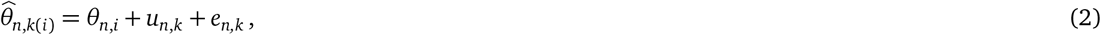

where 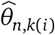 is the estimate for the *n*-th crop growth parameter from Stage 1, *u_n,k_* a spatially correlated random component, and *e_n,k_* are plot residuals assumed to be normally distributed with zero mean and 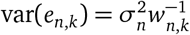, where 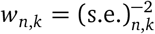 are weights based on the standard error estimates (s.e.) from Stage 1. For a stage-wise approach with weights based on variance estimates of adjusted means, one usually fixes *σ*^2^ to unity (Piepho et al.,2012). Nevertheless, if expecting proportionality of var(*e_k_*) to 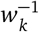 only—for example when the s.e.’s are derived from a correlated trait—it is required to estimate *σ*^2^ as a constant of proportionality. The spatially correlated error term *u_k_* will absorb any spatial correlation caused by random measurement errors and by physical phenotypic differences, and *e_k_* any plot-specific residual.

This approach is not limited to parametric or dimensionality reduction techniques (e.g.,Kwak et al., 2016) but allows including arbitrary dynamic models’ *g* with high complexity based on biologically meaningful traits. Nevertheless, it also obviates modeling a spatio-temporally correlated residual term in its full extent by assuming that all serial correlation is accounted for by the time-dependence of *g*. In the following, we hypothesize and exemplify with a simulation that our approximation of the spatio-temporal correlation structure is well suited to extract intermediate traits with adequate precision from HTFP data.

### 2.2. Dynamic modeling of three trait categories

In dynamic modeling, one has to specify a method, based on *g* of Equation 1, to estimate a vector of meaningful plot-level traits 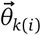 (for brevity we henceforth drop the index *i* for genotypes, referring to 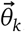, it being understood that a plot-level parameter is always genotype-specific) based on measured phenotypes *y_kt_* and measured covariates *x* at (potentially differing) time points *t*. In the following, we will provide theoretical considerations and specific examples for each of the three trait categories defined in the introduction, (1) timing of key stages’ traits, (2) quantities at defined time points or periods, and (3) dose-response curve traits.

The first intermediate trait category—timing of key stages—describes growth as a sequence of key stages. Such phenology traits are well-known in agronomy, e.g., the timing of jointing (start of stem elongation), heading, and flowering in wheat.

The second intermediate trait category—quantities at defined time points or periods—describes phenotypic characteristics at key stages or steady state phases. Hence, such traits include a time point definition with traits of the first category. The number of tillers per plant at jointing, the number of ears per square unit at harvest, or the average canopy cover between tillering and jointing are examples of such traits for wheat.

The third intermediate trait category—dose-response curves—describes phenotypes as the result of a dose-response model dependent on a covariate course between key stages. Hence, also these traits require time point definitions, e.g., with traits of the first category. The response of the stem elongation to temperature is an example of such a trait for wheat.

Biological drivers of the timing of phenological stages (which are related to all three trait categories) are manifold. Despite this complexity, research in model organism such as *Arabidopsis thaliana* (Wilczek et al., 2009) as well as crops such as wheat (Hyles et al., 2020) has shown that the phenology of outdoor-growing plants can be linked closely to the genetics. In agriculture, such phenological stages are often expressed in thermal time. Thermal time is a widely accepted concept (Parent et al., 2019) and almost 300 years old (Wang, 1960). Still, thermal time is just a mental construct (McMaster and Wilhelm, 1997) that allowed researchers to mask—yet unknown—biological mechanism. Nowadays, gaining insight into those biological mechanisms and manipulating them in desired directions in breeding programs may call for more mental-model-free observational approaches.

Consequently, to obtain traits of the first two categories, we favor semi-parametric approaches that require less biological assumptions (e.g., spline fitting) over parametric approaches (e.g., logistic regression based on thermal time) for the dynamic modeling in Stage 1. When using a semi-parametric approach (such as P-splines), one approximates *g* with a plot-specific model as a smooth function of time *s*(*t*). To extract traits of the first category—timing of key stages—from such a smooth function, a set of methods *q_n_* (*n* = 1,…,*N*) to estimate timing traits *θ*^*T*(*n*)^ (e.g., to approximate the end of the stem elongation phase) from *s* has to be defined,

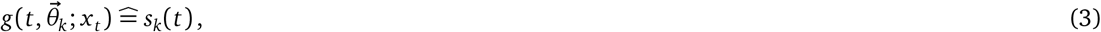

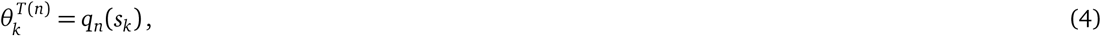

where 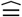 indicates that *s_k_* approximates *g* for the *k*-th plot.

Extracting traits of the second category—quantities at defined time points or periods—builds on the spline function *s* (Equation 3) and extracted timing of key stages (Equation 4) but inverts the approach of extracting key stages: If *θ*^*T*(*n*)^ represent timing of key stages (e.g., the end of stem elongation), then quantities at defined time points *θ*^*Q*(*n*)^ (e.g., canopy cover at the approximated end of stem elongation) may be extracted from the spline *s* as

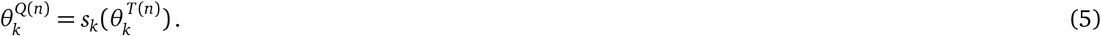

It is important to note that the underlying low-level traits for the timing trait *θ*^*T*(*n*)^ and the spline *s* in Equation 5 may differ, giving rise to a vast amount of possible trait combinations, such as combining canopy height timing traits with canopy cover quantity traits. While Equation 5 extracts quantities at points in time, extracting aggregated quantities (e.g., normalized area-under-the-curve traits) for a period of time may be of interest as well. If *θ*^*T*(*a*)^ and *θ*^*T*(*b*)^ represent two cautiously chosen timings of key stages’ traits where *θ*^*T*(*a*)^ < *θ*^*T*(*b*)^ (e.g., approximated start and end of flowering), then a quantity at defined time period trait *θ*^*Q*(*a…b*)^ (e.g., average temperature at approximated flowering) may be extracted from *s* as

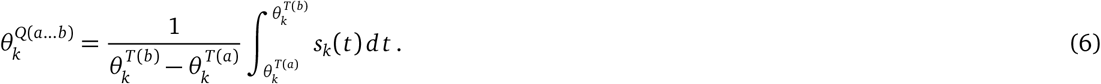

If either 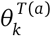 or 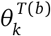 corresponds to a time series boundary (e.g., end of stem elongation to end of time series), the trait may represent an initial or final trait value (e.g., final height).

For the third trait category—dose-response curves—one describes a phenotype as the result of a dose-response model *ġ* that relates growth rates to a covariate course *x_t_* and a corresponding set of crop growth model parameters *θ^C^* = (*θ*^*C*(1)^, *θ*^*C*(2)^,…, *θ*^*C*(*L*)^) where *L* is the total number of parameters of the dose-response curve,

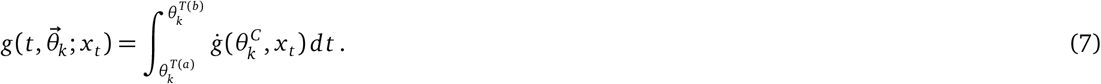

Similar to quantities at defined time periods’ traits (Equation 6), dose-response curve traits require the definition of a corresponding growth phase, characterized by a start (*θ*^*T*(*a*)^) and a stop (*θ*^*T*(*b*)^). Therefore, a preliminary extraction of traits of the category one (Equation 4) is required. Subsequently, *θ^C^* may be estimated.

The striking similarity of Equation 6 and 7 is no coincidence. The area-under-the-curve of a defined growth period can be seen as a direct cause of a response to covariates in this growth phase. The two approaches differ in how they include covariates: While dose-response curves model an explicit dependency on covariates, an area-under-the-curve quantifies implicitly the result of such a dependency.

An example for a dose-response curve *ġ* at a defined growth phase is the stem elongation rate of wheat in relation to temperature. Extracting such a dose-response curve implies that one is interested in fitting a specific non-linear function.

### 2.3. Combining multi-year measurements

HTFP platforms such as the FIP are usually run on a continuous basis, thus increasing the number of year measurements with each year of operation since inauguration. Experimental designs and genotype sets may change to some extent across the years. The question is how to combine such multi-year measurements in a way that one can process years in stages, which is of major benefit for both documentation purpose and processing requirements.

The problem of stage-wise analysis we are addressing here has a long history (Cochran, 1954) and is well known in plant breeding (Smith et al., 2005;Piepho et al., 2012) and also in other contexts, most notably in meta-analysis (Whitehead, 2002;Borenstein et al., 2009). Most commonly, the problem arises in settings where information needs to be combined across several experiments, whereas in the present work we consider the case where different pieces of information need to be combined across units in a single experiment. Despite these differences in scale, the statistical challenges are the same. To illustrate, consider a simple setting in which a set of replicated genotypes is tested for yield over a number of years in a platform. The response of the *i*-th genotype on the *k*-th plot at the *j*-th year can be written as

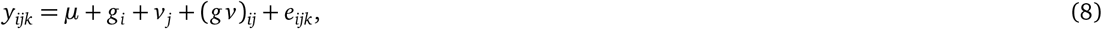

where *μ* is an intercept, *g_i_* is the main effect of the *i*-th genotype, *v_j_* the main effect of the *j*-th year, assumed to be normal with zero mean and variance 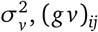 is the interaction of the *i*-th genotype and *j*-th year assumed to be normal with zero mean and variance 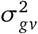, and *e_ijk_* a residual error assumed to be normal with mean zero and year-specific variance 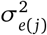. Note that depending on the environments to examine, more complex G×E models than the one introduced in Equation 8 ((*gv*)_*ij*_) may be preferable (van Eeuwijk et al., 2016). An objective among others in field phenotyping platforms is to estimate genotype means across years, *η_i_* = *μ*+*g_i_* and their differences.

This can be done in a single stage by fitting the model (Equation 8) directly to plot data *y_ijk_*. Alternatively, we may proceed in two stages and first estimate genotype means per year using sample means 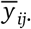. These means have variance 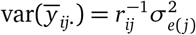, where *r_ij_* is the number of replications of the *i*-th genotype in the *j*-th year. In the second stage, we can fit the model

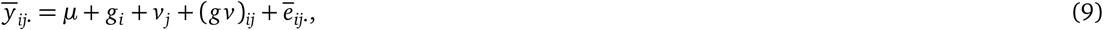

where 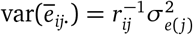, which is the conditional variance of the genotype means computed in the first stage. The estimates of genotype means, *η_i_* = *μ*+*g_i_*, are identical for single-stage and two-stage analysis, provided the variance components are known (Piepho et al., 2012). Differences arise in practice because variances need to be estimated. Stage-wise analysis entails an approximation of the gold standard of single-stage analysis because variances 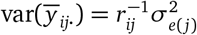 as estimated in the first stage are treated as known quantities in the second stage, disregarding the degrees of freedom associated with these estimates and their uncertainty. A key feature of stage-wise analysis is that the inverses of these estimated variances act as weights in the second-stage analysis. A major challenge in any stage-wise analysis is how to best determine the weights and how to account for the uncertainties associated with them.

The situation faced in the analysis of HTFP is comparable in that it proceeds in stages with necessity because a single-stage analysis is in conflict with performance and generalization demands (i.e., multi-year HTFP data may comprise a number of differing experimental designs that require individual processing in a first stage) and that the primary interest is the genotype main effect *g_i_*, which equals *θ_i_* in HTFP (Figure 3k–m). The statistical challenges are rather more daunting, however, for several reasons: (i) HTFP involves high-frequency time series in which observations are serially correlated; (ii) summarizing time-series data usually requires nonlinear regression models; (iii) analyses of field trials are often done exploiting spatial correlations among neighboring plots; (iv) remote or proximate sensed data are affected by environmental conditions (wind, illumination) that may change during the course of a measurement; (v) the number of processing steps required for the full analysis process is much greater than two (see the number of transitions between stages in Figure 3 (grey arrows)). These additional features make the determination of appropriate weights to be carried forward from one stage to the next even more challenging than in the simple example given above.

Here, we propose a weighting approach for the intermediate trait category (1) (timing of key stages) and (2) (quantities at defined time points or periods) only for brevity, and illustrate its application using a simulation study described in the following section. Traits of the third category (dose-response curves) will be considered in a follow-up paper (Roth et al., 2021).

### 2.4. Simulation of canopy height data

To demonstrate the extraction of traits of the first two categories (timing of key stages and quantities at defined time point or periods), winter wheat canopy height data were simulated implementing a temperature dose-response curve (trait category three, Equation 7). The temperature response of the stem elongation phase was assumed to follow a dose-response curve with break points (Figure 4),

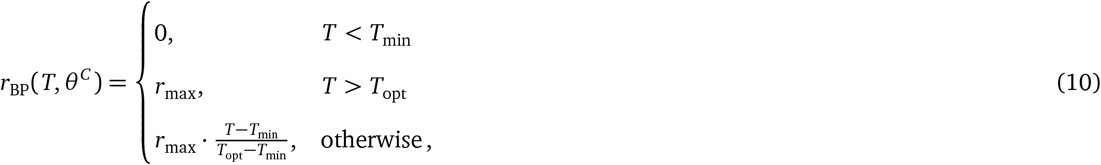

where *T*_min_ is the base temperature below which the elongation rate *r* is zero and *T*_opt_ the optimum temperature above which the elongation rate reaches the maximum hourly elongation rate *r*_max_, while *θ^C^* = (*r*_max_, *T*_min_, *T*_opt_) (Figure 4).

**Figure 4:**
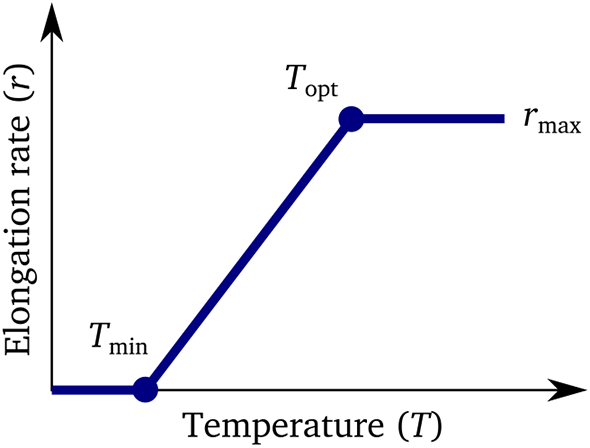
Schematic drawing of the dose-response model (*ġ* of Equation 7) implemented as break-point model (*r*_BP_, Equation 10) used for the simulation of canopy height time series based on temperature courses.

As starting point for the simulation, existing experimental designs of three consecutive years at the ETH research station of agricultural sciences in Lindau Eschikon, Switzerland (47.449 N, 8.682 E, 556 m a.s.l.) were used. The experiment consisted of 352 wheat genotypes, replicated twice per year on two spatially separated fields, both augmented with spatial checks in a 3×3 block arrangement.

To simulate canopy height time series, existing weather data were used to introduce a close-to-realistic stochastic behavior. Canopy growth was simulated for a measurement interval of one per day and for a period between first of March and 20th of July (*d* = 1 ≤ *t* ≤ *d* = 142) for each of the three simulated years *j* (*j* = 2016,2017,2018). Growth between daily campaign time points *t* was modeled as cumulative response to hourly temperature measurements *T_jdh_* (*h* = 1,…,24). The canopy height *y_ijkt_* of genotype *i* (*i* = 1,…,352) at plot *k* (*k* = 1,…, 704) in the year *j* at a specific time point (*t* = 1,…, 142) was then simulated as

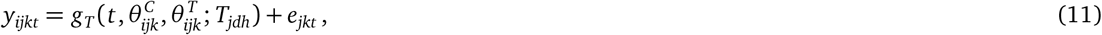

where *g_T_* depends on *r*_BP_ in Equation 10 (see below) and simulates growth as a function of temperature *T_jdh_*, time point *t*, a vector of plot-specific crop growth model parameters 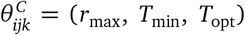, and a vector of plot-specific timing traits 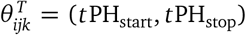. The error term *e_jkt_* simulates plot and time point residuals.

The growth function *g_T_* was specified as

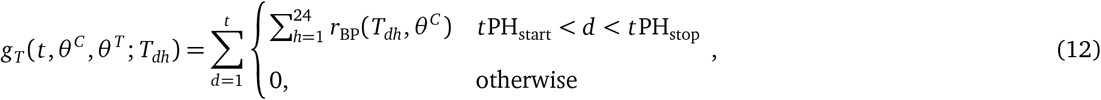

where *r*_BP_ represents a dose-response as function of hourly temperatures *T_dh_* and a vector of crop growth model parameters *θ^C^* (Equation 10), *t*PH_start_ the time point where canopy growth started, and *t*PH_stop_ the time point where canopy growth stopped.

This approach produced realistic-looking canopy growth curves (compare Figure 5 with, e.g., real data in Kronenberg et al. 2017, 2020a) with a characteristic start of growth (*t*PH_start_) and a stop of growth (*t*PH_stop_), corresponding to the first intermediate trait category (timing of key stages). Additionally, growth curves indicated a characteristic final height (PH_max_), corresponding to the second intermediate trait category (quantities at defined time points or periods).

**Figure 5:**
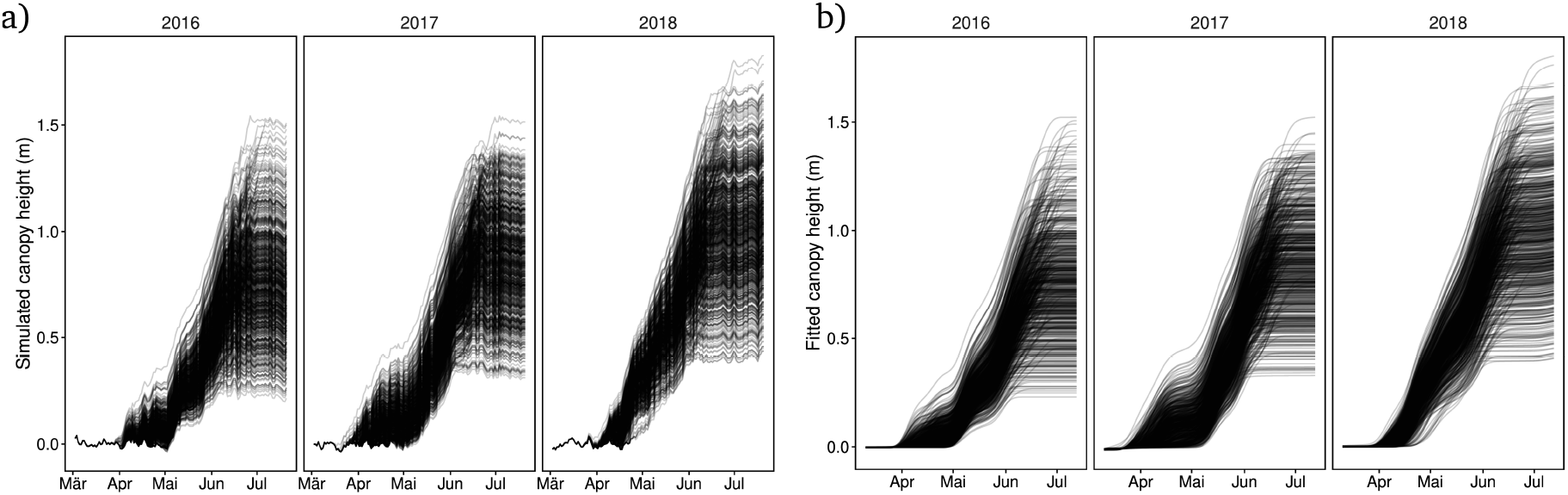
Simulated canopy heights (a) and fitted canopy height splines (b) for one simulation run with 352 genotypes, two replications per year, and three years, corresponding to the proposed 3-stage temporal-first (*t*→*k*→*j*) approach.

Noise as specified in Section 2.1 was introduced on a genotype, plot and time point level. Genotype-year interactions were not explicitly introduced but assumed to emerge from an intrinsic property of the simulation. The simulation was based on applying a dose-response curve to temperature courses. Therefore, the combination of the genotype-specific parameters 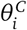 and 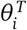 and year-specific temperature courses will lead to differently shaped growth curves for one genotype in different years. Those differences may then emerge as rank shifts of simulated canopy heights at specific time points, i.e., G×E.

To add noise to genotype traits, the crop growth model parameters 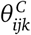 and the timing traits 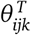 were further decomposed in genotypic and spatially correlated parts,

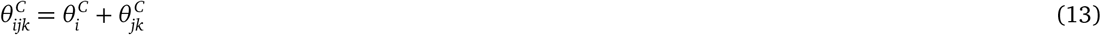

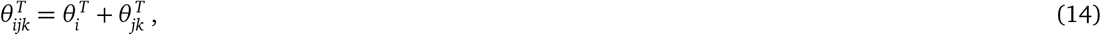

where 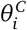 and 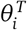 were simulated using normal distributions 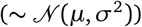. Trait-specific *μ* and *σ*^2^ were chosen based on literature if available, and otherwise based on own unpublished field data. 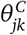 and 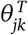 were spatial correlated heterogeneity components for those traits (AR(1)_x_⊗AR(1)_y_), where AR(1)⊗AR(1) is a two-dimensional first-order autoregressive model in row (*x*) and range (*y*) direction, mimicking the influence of other covariates and therefore spatial heterogeneity. Note that a high autocorrelation in row and range direction with *ρ*_*x*&*y*_ = 0.95 and half the variance of the corresponding input parameter (Appendix, Section 5.1, Table 2) was assumed, which appeared reasonable for cereal experiments (Patterson and Hunter, 1983;Velazco et al., 2017) when considering that the plot residual components will dilute the marginal correlation of the whole residual structure.

The plot residual *e_jkt_* was simulated as sum of three error terms,

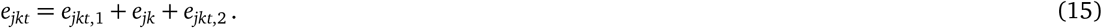

The first error term *e*_*jkt*,1_ corresponds to the serial correlation of measurement errors (AR(1)_t_) that *g* in Equation 1 presumably absorbs. The second error term *e_jk_* mimics a systematic spatially correlated measurement error after an incomplete correction with reference measurements (AR(1)_x_ ⊗AR(1)_y_). We note that adding this error introduces an intentional discrepancy between the analysis model and the simulation: the proposed plot-level model for repeated measurements does not include such a systematic error in the first stage (dynamic modeling). Consequently, estimating the spatial correlation in the second stage will confound measurement errors and nuisance factors, which corresponds to a situation we frequently encounter in HTFP. The third error term *e*_*jkt*,2_ corresponds to *e_kt_* in Equation 1 and represents a plot-based i.i.d. residual 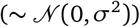. The first error term was assumed to cause most of the known measurement error, wherefore the corresponding *σ* (*σ_m_*) was set accordingly to 0.01 m (Roth et al., 2020), but significantly reduced for the second error term (*σ*_*ex*&*y*_) and third error term (*σ_e,k_*). The autocorrelation parameters *ρ* for the first (*ρ_m_*) and second (*ρ*_*ex*&*y*_) error term were arbitrary set to 0.7. All simulation input parameters and sources for the aforementioned assumptions are summarized in the Appendix (Section 5.1, Table 2).

A total of 500 simulation runs were performed. These simulated time series with a measurement interval of one day were then further thinned to intervals of three, five, seven and 14 days to study the effect of lower frequencies.

We note that the simulation (Equation 11) comprised *θ^T^*, i.e. traits of the first category, and *θ^C^*, i.e. traits of the third category. The second trait category *θ^Q^* was dependent on the first and third category and year specific temperature courses, and therefore only an indirect input parameter of the simulation. Therefore, the simulation allowed extracting traits of all three categories, and validating traits of category one (*θ^T^*) and three (*θ^C^*) with genotypic input data, and traits of category two (*θ^Q^*) with plot-level (indirect) input data. Here, we illustrate the extraction of *θ^T^* and *θ^Q^* only for brevity. The extraction of *θ^C^* and therefore dose-response curve parameters of a crop growth model will be considered in a follow-up paper (Roth et al., 2021).

We further note that all simulation input parameters for a given genotype *i* in 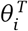 and in 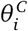 were uncorrelated, for *n* in 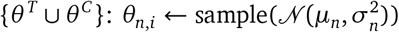.

In reality, genetic effects and artificial selection have certainly resulted in weak to strong correlations for those parameters. Dynamic modeling may introduce new, artificial correlations of parameters. An example for such an artificial correlation is the dependency of the measurable start of the stem elongation on the timing parameter *t*PH_start_ and the dose-response parameter *T*_min_. Extreme values in those parameters may lead to growth curves with a comparably delayed measurable start of the stem elongation phase. Consequently, although genetically uncorrelated, the extracted parameters will be correlated. When examining a real-world genotype set, e.g., a breeding population, these effects will be confounded, but using a simulation with uncorrelated input parameters allows quantifying the extraction artifacts.

### 2.5. Stage 1: Extracting the timing of key stages and quantities at defined time points

To extract timing of key stages, a monotonically increasing P-spline was fitted to plot time series using the R package *scam* (Pya, 2019), thus implementing *s_k_*(*t*) of Equation 3. The package fits shape constrained generalized additive models (GAM) (Pya and Wood, 2015). A Bayesian approach to uncertainty quantification is used to obtain standard errors of predictions.

The number of knots was set proportional to three quarters of the (thinned) observations. In a next step, the start and end of stem elongation (*t*PH_start_ and *t*PH_stop_) were extracted based on the quarter of maximum elongation rate (QMER) method, which in brief extracts key time points with elongation rates greater than a threshold of one quarter of the maximum elongation rate. Thus, the QMER method represents an implementation of *q_n_*(*s_k_*) of Equation 4.

The decision to chose one quarter of the maximum elongation rate is reasoned as follows: The maximum elongation rate of wheat stems according to our own data is around 1 mm h ^-1^ (Table 1). Given a measurement interval of one to 14 days and the number of spline knots set to three-quarter of the number of measurements, this will result in height differences between spline knots of approximately 30 mm to 450 mm for the maximum elongation rate, and 10 mm to 150 mm for one third of the maximum elongation rate. The measurement error for canopy height measurements is known to range around 10 mm (Table 1). Consequently, the measurable differences between estimates at spline knots are larger than the measurement error if growth rates are greater than one third of the maximum elongation rate, which should allow for a detection of turning points that represent a transition from growth rates close to zero to growth rates close to the maximum and *vice versa*.

**Table 1:** Model parameters for the second and third stage of the stage-wise linear mixed model analysis. *k* denotes the *k*-th plot, *j* the *j*-th year, and *i* the *i*-th genotype.

| Stage | Term | Description | Part |
| --- | --- | --- | --- |
| 2) | $\hat{\theta}_{jk}$ | Plot response based on dynamic modeling | Response |
| | $\theta_{ij}$ | Year genotype response | Fixed |
| | $p_{c(jk)}$ | Range effect on field (main working direction, e.g., for sowing) | Random |
| | $p_{r(jk)}$ | Row effect on field (orthogonal to main working direction) | Random |
| | $f_n(x(jk), y(jk))$ | Smooth bivariate surface in spatial $x$ and $y$ coordinates (mapping real distances on the field) consisting of a bivariate polynomial and a smooth part (for details see Rodríguez-Álvarez et al., 2018) | Spatial |
| a) | $e_{jk}$ | Residuals with $\text{var}(e) = \sigma^2$ | Residual |
| b) | $e_{jk}$ | Residuals with $\text{var}(e) = \sigma^2 w^{-1}$, where $w$ are weights based on the standard error estimates from the previous dynamic modeling step (Stage 1), and $\sigma^2$ the residual variance parameter | Weights |
| 3) | $\hat{\theta}_{ij}$ | Adjusted year genotype mean (BLUE) from Stage 2 | Response |
| | $\mu$ | Global intercept | Fixed |
| | $\nu_j$ | Year effect | Random |
| | $\theta_i$ | Genotype response | Fixed |
| | $(\theta_n \nu)_{ij}$ | Genotype year interaction | Residual |
| a) | $e_{ij}$ | Residuals with $\text{var}(e) = \sigma^2$ | Residual |
| b) | $e_{ij}$ | Residuals with $\text{var}(e) = \sigma^2 w^{-1}$, where $w$ are weights based on the square rooted diagonal of the variance-covariance matrix from Stage 2, and $\sigma^2$ the residual variance parameter | Weights |

In the first step, spline predictions for canopy heights 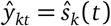 and standard error estimates s.e.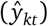 were calculated separately for each plot at hourly time steps using the prediction function of the *scam* package. Thereafter, hourly growth rates 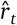 were derived from the difference between subsequent predictions, 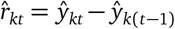 (Figure 3e). Then, the following algorithm was applied to extract intermediate traits and corresponding weights *w* based on standard errors of spline predictions for each plot (*k* is omitted in the following for sake of simplicity):

1. Determine maximum elongation rate:

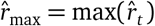
2. Filter 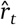 for data points with an elongation rate greater than ¼ of the maximum elongation rate:

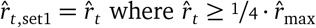
3. Define the earliest time points that are left after filtering as the start of growth:

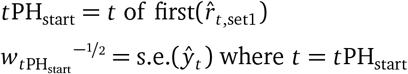
4. Filter 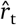 for data points with an elongation rate lower than ¼ of the maximum elongation rate and a minimum distance of 40 days to the approximated start of growth:

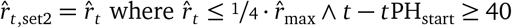
5. The earliest value that is left after filtering indicates the approximated end of growth:

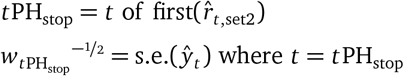

Note that the weights for timing of key stages’ traits in this work were based on the standard errors of spline predictions 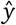. We will address the conditions that should be met to justify our approach in the following section.

We extracted the growth stages start and end of stem elongation (*t*PH_start_ and *t*PH_stop_) and corresponding standard error estimates based on the quarter of maximum elongation rate (QMER) method. To compare the QMER method with the approach taken by Kronenberg et al. (2017), we additionally determined the time points where 15% (*t*PH_15_) and 95% (*t*PH_95_) of final height was reached (for details, see Kronenberg et al., 2017). In Figure 3e, we depict only the QMER method.

The quantity at a defined time point final height (PH_max_) was calculated as the median of 24 hourly spline predictions after the estimated stop of growth:

1. Filter 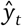 for data points after reaching final height:

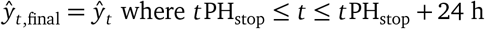
2. Aggregate data points:

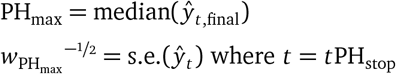

### 2.6. Weigting based on estimated standard errors

The chosen implementation of the QMER method did not provide standard errors for the derived growth rate 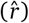 and time points (*t*). Instead, weights for further processing after the dynamic modeling were based on standard errors of spline-based predictions of the response (s.e.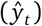). Using weights based on the standard errors of spline predictions is intuitive for quantities at defined time points or periods’ traits (e.g., PH_max_), as both s.e.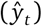 and 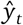 share the same unit. However, for timing of key stages (e.g., *t*PH_start_ and *t*PH_stop_), such a weighting approach requires a positive and high association between the true weights for *t* and *y* for a given (to be determined) time point. Alternatively, one could use an inverse regression approach (e.g., the Fieller’s theorem (Seber, 2003) or the delta method (Johnson et al., 1993)) to determine weights for two means with different units. Such an inverse regression approach becomes non-trivial when involving a combination of statistical tools—e.g., P-splines and the QMER method. Therefore, using an inverse regression approach may contradict the requirement to provide a seamless workflow to integrate arbitrary complex dynamic models *g* (Equation 2).

Consequently, we decided to assume proportionality of weights for standard errors of spline predictions and timing of key stage estimations. The factor of proportionality was estimated via the residual variance (*σ*^2^), which was estimated in each analysis, rather than fixed at unity, as is customary in standard weighted analysis, where the inverse weights are taken to be the known residual variances (Piepho et al., 2012). Our assumption is based on considerations on a concrete example (see Appendix, Section 5.3). In addition, standard errors of spline predictions suppose that observations of plot-based time series are independent. As this is—at least for the simulation—not true (see Section 2.1), the calculated standard errors of the estimates will be biased. To test whether weighting was advantageous, despite possible bias in the weights and imperfect proportionality for timing of key stage traits, we optionally provided these weights in the next processing step.

### 2.7. Stage 2: Calculating adjusted genotype means per year

The extraction of dynamics characteristics resulted in measurement time point independent trait values at a plot level (Stage 1). These plot values were subsequently processed in a two-stage linear mixed model analysis (Stage 2 and 3), where the second-stage analysis averaged over within-year effects (e.g., spatial heterogeneity) and the third-stage analysis over between-year effects.

We used SpATS (Rodríguez-Álvarez et al., 2018) to fit a model with a smooth bivariate surface defined over spatial coordinates of plot centers (*f* (*x*(*jk*), *y*(*jk*))) and added fixed genotype effects (*θ_ij_*) and random effects of plot rows and ranges (*p*_*r*(*jk*)_ and *p*_*c*(*jk*)_),

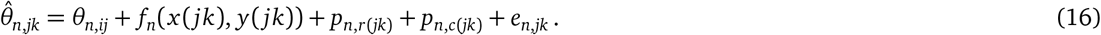

Model parameters are listed and explained in Table 1 (Stage 2). Stage 2a and 2b are two nested models; Stage 2b corresponds to Stage 2a but additionally includes weights. Equation 16 was applied to all intermediate traits to calculate BLUEs of genotype means per year.

### 2.8. Stage 3: Genotypic marginal means calculation

The second stage already covered aspects such as spatial heterogeneity and design-specific characteristics such as row and range arrangements, and allowed obtaining adjusted year genotype means 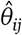 (BLUEs). In the third stage, those means were further processed with a model based on Equation 9,

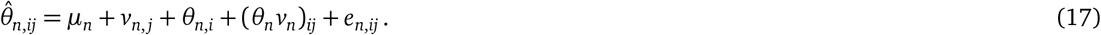

The model assumes that genotype-environment effects can be partitioned into genotype response effects (*θ_i_*) and genotype-year interaction effects ((*θv*)_*ij*_)(Piepho et al., 2012) while the residual errors (*e_ij_*) are assumed to be identically and independently normally distributed. Model parameters are listed and explained in Table 1 (Stage 3). Stage 3a and 3b are two nested models; Stage 3b corresponds to Stage 3a but additionally includes weights. Models were fitted using the R package ASReml-R (Butler, 2018). Equation 17 was applied to all intermediate traits to calculate overall genotype BLUEs.

### 2.9. Comparison with a two-stage approach

Separating the dynamic modeling step (*t*) from further processing steps (*kj*) prevents implementing the gold standard of a one-stage analysis. Nevertheless, subsequent processing stages can be summarized in one stage, hence resulting in a two-stage temporal-first approach (*t*→*kj*). To allow comparing such an approach with the proposed three-stage approach (*t*→*k*→*j*), the estimated intermediate traits from Stage 1 were additionally processed using a two-stage model,

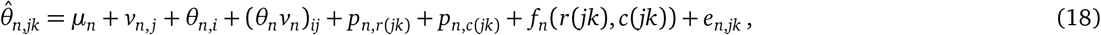

where *μ* is a global intercept, *v_j_* a year intercept, *θ_i_* the genotype response, (*θv*)_*ij*_ genotype year interactions, *p*_*r*(*jk*)_ and *p*_*c*(*jk*)_ range and row effects, *f* (*r*(*jk*), *c*(*jk*)) year specific AR(1) ⊗ AR(1) interactions based on ranges (*c*()) and rows (*r*()) of plots, and *e_jk_* plot residuals with year-specific variances.

### 2.10. Comparison with a three-stage spatial-first approach

In this work, we presented a strategy to process HTFP data, starting with dynamic modeling (*t*), followed by two stages of a linear mixed model analysis, first averaging over within-year effects (*k*) followed by averaging over between-year effects (*j*), thus, *t*→*k*→*j*. This approach is to some extent the reverse of van Eeuwijk et al. (2019) who suggested correcting time point measurements in a first stage of a stage-wise linear mixed model analysis, followed by dynamic modeling and modeling of environmental dependencies, and a second stage of a stage-wise linear mixed model analysis to calculate adjusted means across years, thus, *k*→*t*→*j*.

To allow a comparison with the strategy van Eeuwijk et al. (2019) proposed for the examined timing of key stages traits, we additionally implemented the *k*→*t*→*j* approach. To do so, in a first step, Equation 16 was applied to low-level trait measurements per time point *t* using SpATS (Rodríguez-Álvarez et al., 2018),

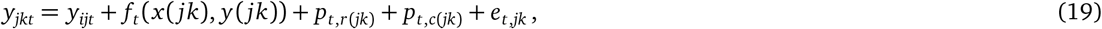

This step resulted in estimated means and standard errors per time point for *y_ijt_*. Then, dynamic modeling (Equation 1) was applied to these genotype time point estimates 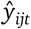 using P-splines and the QMER method as described in Section 2.5,

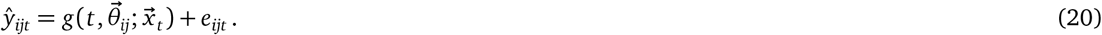

The residual variance was set to var(*e*) = *σ*^2^*w*^−1^ where *w* are weights based on the standard error estimates from the previous spatial modeling step, and *σ*^2^ is the residual variance parameter. This step resulted in estimates and standard errors for the crop growth model parameters 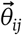 at the genotype level. For each of these estimated parameters 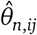, in a last step, Equation 17 was applied to calculate genotypic marginal means,

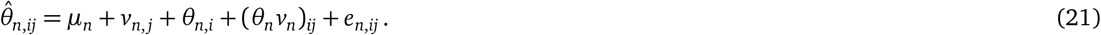

The residual variance was set to var(*e*) = *σ*^2^*w*^−1^ where *w* are weights based on the standard error estimates from the previous dynamic modeling step, and *σ*^2^ is the residual variance parameter. Consequently, we applied a fully weighted spatial-first approach (*k*→*t*→*j*) to the simulation.

### 2.11. Simulation validation

Bias, variance, root-mean squared error (RMSE) and Pearson’s correlation were calculated both after dynamic modeling (Stage 1) and after the stage-wise linear mixed model analysis (Stage 2 and 3) separately for each simulation run.

## 3. Results

A total of 176,000 genotype-runs (352 genotypes × 500 runs) replicated on 1,056,000 plots (number of genotype runs × 3 years × 2 replications) containing 149,952,000 data points (number of plots × 142 measurement days) were simulated. In the following, we give insights on the precision of extracted traits influenced by the choice of method, weighting, and measurement interval.

### 3.1. Dynamic modeling

P-splines model fits converged for all simulated plot time series and produced smooth-looking growth curves (Figure 5). Start and end of stem elongation estimations were successfully extracted using the QMER method as well as the final height percentile method.

The timing of the key stage trait *t*PH_start_ was better estimated by the P-spline/QMER method with a lower median RMSE and lower median bias (Figure 6). Nevertheless, in comparison to the final height percentile method, the median variance was higher, and larger outliers for RMSE and variance were found. The trait *t*PH_stop_ was better estimated by the final height percentiles method with lower median bias, median RMSE and median variance than by the P-spline/QMER method, but the percentiles method also produced larger outliers for variance and RMSE than the P-spline/QMER method.

**Figure 6:**
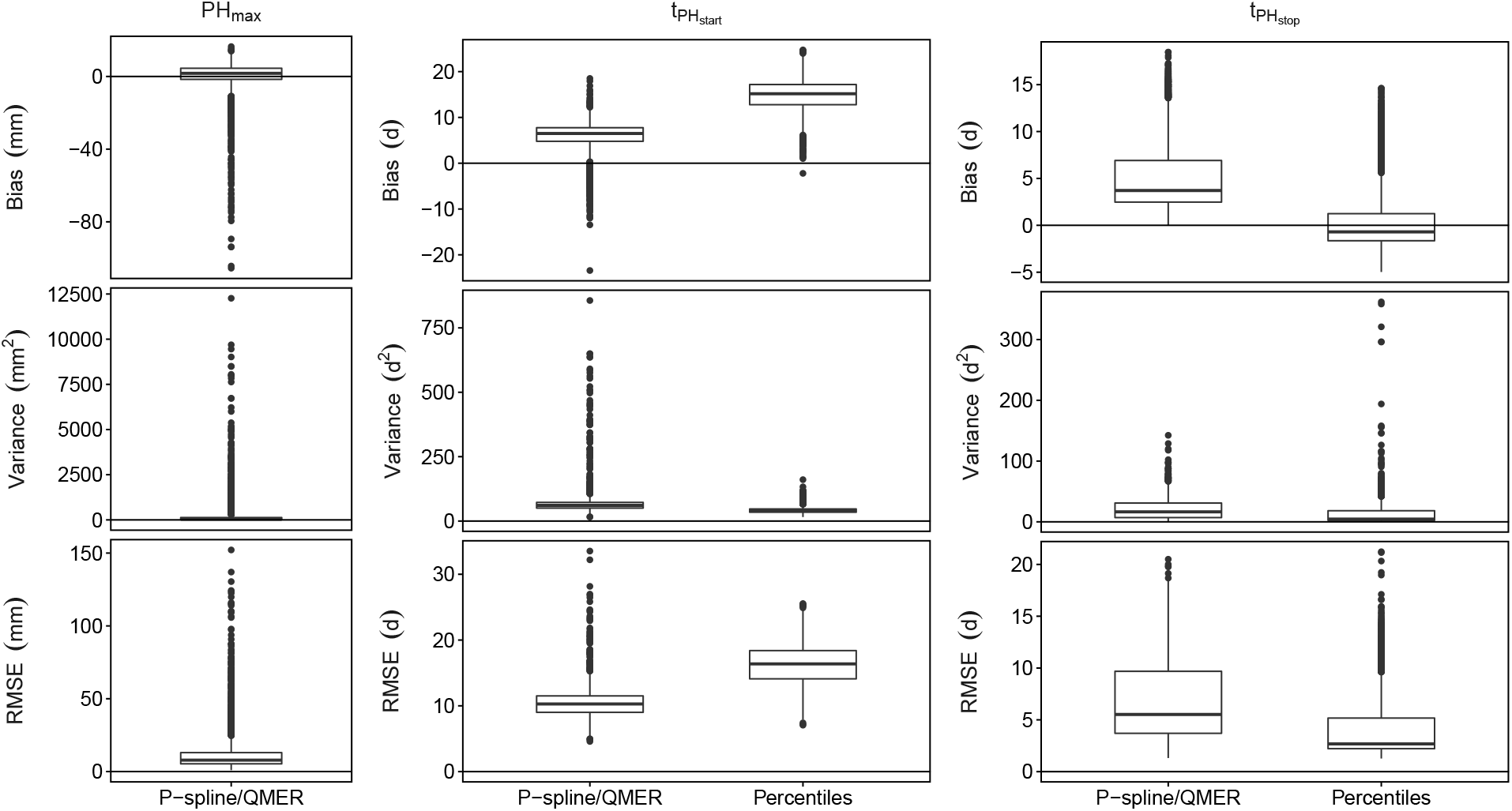
Box plots for the 500 simulated datasets of plot-based bias, variance and root-mean squared error (RMSE) of two timing of key stages models (P-spline/QMER model and final height percentiles), corresponding to the proposed 3-stage temporal-first (*t*→*k*→*j*) approach.

Both the P-spline/QMER and final height percentile methods performed comparably and were able to predict *t*PH_start_ with a strong and *t*PH_stop_ with a very strong correlation to input values (Figure 7), but also for both methods, the estimated start of stem elongation (*t*PH_start_) was weakly biased by the input trait base temperature. Nevertheless, the correlation between the extracted start and end of stem elongation—an artifact of the method, as the simulation input was uncorrelated—was much higher for the Percentile method than for the P-spline/QMER method. Based on these findings, the P-spline/QMER model was selected for further processing in the stage-wise analysis.

**Figure 7:**
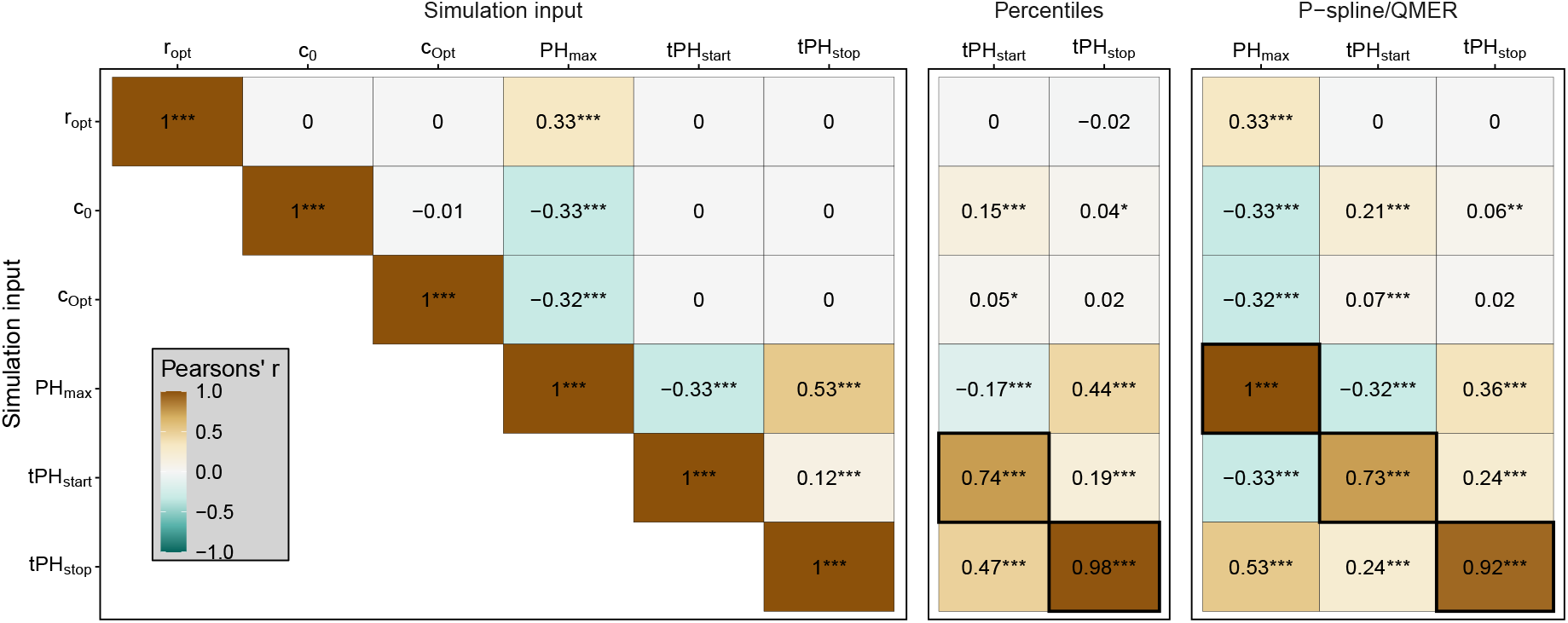
Pearson’s correlations of plot time series traits, corresponding to the proposed 3-stage temporal-first (*t*→*k*→*j*) approach. Provided are simulated input parameters and extracted timing of key stages’ and quantities’ traits for the P-spline/QMER and final height percentile model. On the two right panels, black bold boxes indicate correlations between predicted and true values for identical traits, while all other boxes indicate correlations that arose as artifacts of the extraction. Note that input parameters were uncorrelated (left panel), except for PH_max_.

### 3.2. Required measurement intervals

Estimating *t*PH_stop_ and PH_max_ using the P-spline/QMER or Percentile method was not affected by increased or reduced measurement intervals unless reduced from 7 to 14 days, where the correlation for both *t*PH_start_ and *t*PH_stop_ dropped (Figure 8). The estimation of *t*PH_start_ was, in contrast to the two other traits, sensitive to reduced measurement intervals above five days for the P-spline/QMER method. The prediction of final height was not affected by increased measurement intervals.

**Figure 8:**
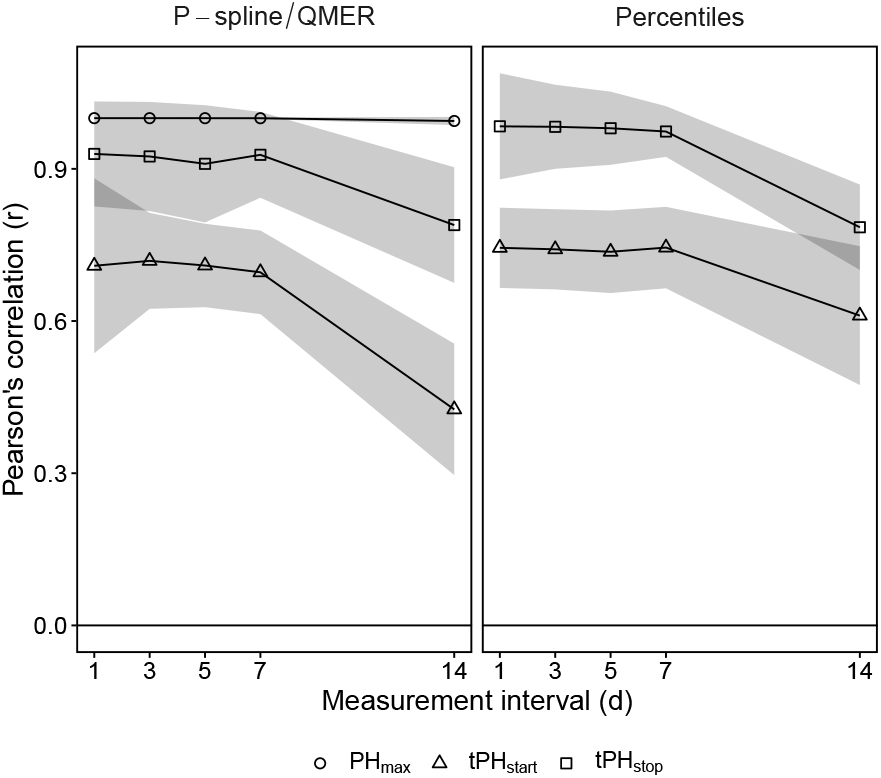
Pearson’s correlations for differing measurement intervals for the timing of key stages based on splines (P-spline/QMER method) and final height percentiles (Percentiles method), corresponding to the proposed 3-stage temporal-first (*t*→*k*→*j*) approach.

### 3.3. Stage-wise linear mixed model analysis

For both traits *t*PH_start_ and *t*PH_stop_, calculating overall adjusted genotype means reduced the median variance and median bias if compared to plot-based values for the P-spline/QMER method (Figure 6) and improved the median RMSE for *t*PH_start_ but not for *t*PH_stop_ (Figure 9, Appendix Section 5.2 Table 3). Based on variance and bias, weighting Stage 2 and 3 with errors of the prediction from the preceding stages was of advantage for *t*PH_start_ and *t*PH_stop_. Nevertheless, for *t*PH_start_ the lowest RMSE with the lowest bias but highest variance was found for the combination of not weighting Stage 2 and Stage 3 (Figure 9, Appendix Section 5.2 Table 3), but differences to weighting both Stage 2 and 3 were very small. In opposite, for *t*PH_stop_, the lowest RMSE with the lowest bias was found for the combination of weighting Stage 2 and Stage 3.

**Figure 9:**
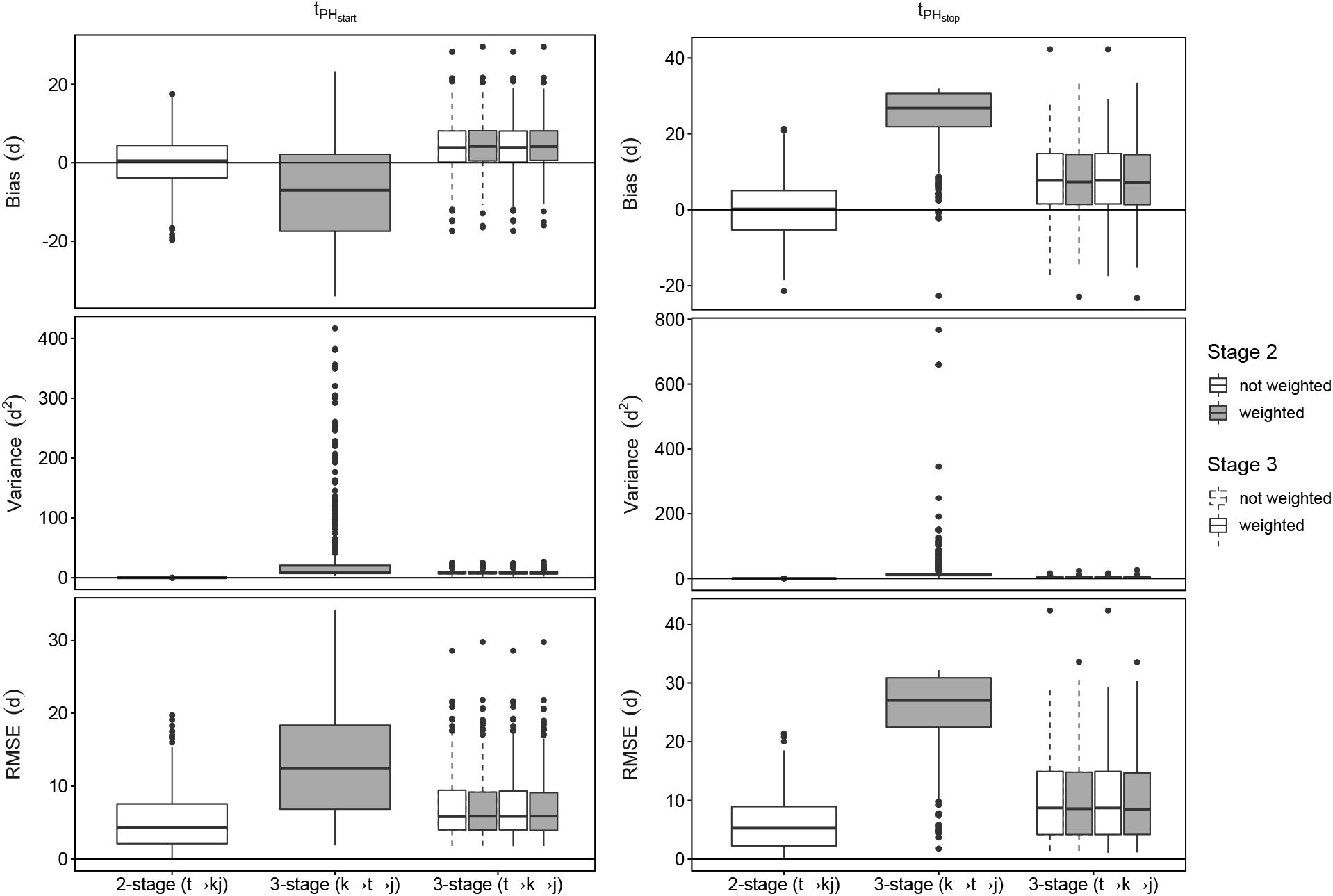
Box plots for the 500 simulated datasets of genotype based bias, variance, and root-mean squared error (RMSE) for the key stages P-spline/QMER model for the spatial-first (*k*→*t*→*j*) and the temporal-first (*t*→*k*→*j*) three-stage model and the temporal-first (*t*→*kj*)two-stage model.

When comparing the proposed three-stage temporal first model (*t*→*k*→*j*) with a two-stage model (*t*→*kj*), using a two-stage model was of advantage for both *t*PH_start_ and *t*PH_stop_, indicated by a lower median RMSE and a higher correlation (Appendix Section 5.2 Table 3) and fewer outliers (Figure 9). Overall, the proposed three-stage temporal-first and the two-stage approach performed comparably, but for the three-stage spatial-first model proposed by van Eeuwijk et al. (2019) (*k*→*t*→*j*), larger differences were found: for *t*PH_start_, while having a very low bias, the spatial-first model resulted in a higher variance and consequently RMSE and a low correlation. For *t*PH_stop_, the bias as well as the variance for the spatial-first approach were very large, resulting in a larger RMSE and a smaller correlation than for the proposed temporal-first approach.

## 4. Discussion

### 4.1. Data processing in stages

The overall workflow of HTFP requires a joint effort of disciplines (Cobb et al., 2013;Araus and Cairns, 2014) which may be separated into three main domains: (1) automation and sensing including feature extraction from sensor readouts, (2) applied phenotyping including dynamic modeling and trait extraction from sensor-derived features, and (3) analysis of designed agricultural experiments or breeding experiments. Plant phenomics must bridge these three disciplines with the overall aim to characterize phenotypes as the result of genotype, environment and management. A plot-level model for repeated measurements may help to link the highly specific domains of sensing and the analysis of experiments. The link to genomic information in breeding and quantitative genetics further increases the complexity of the topic, but is not addressed in this study.

Here, we presented a strategy to process HTFP data. Based on the evaluated sources of variation, we decided to process in stages, starting with dynamic modeling, followed by two stages of a linear mixed model analysis (for a concrete application see, e.g.,Anderegg et al., 2020). This approach is to some extent the reverse of van Eeuwijk et al. (2019) who suggested correcting time point measurements in a first stage of a stage-wise linear mixed model analysis, followed by dynamic modeling and modeling of environmental dependencies, and a second stage of a stage-wise linear mixed model analysis to calculate adjusted means across years. Both options—correcting for spatial or temporal correlations first—represent valid alternatives. Nevertheless, for timing of key stages traits, this study revealed differences in the effectivity of the methods, resulting in larger bias and lower correlations for the spatial-first approach. Compared to the spatially correlated phenotypic variation in the timing of key stages 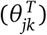, the spatially correlated measurement error (*e_jk_*) was much smaller in our simulation (Appendix, Section 5.1, Table 2). Given this constellation and the indication that a temporal-first approach is closer to the data generating mechanism than a spatial-first approach (for further elaborations and an additional example of such effects, please see the Appendix, Section 5.4), using a temporal-first approach may be of advantage if aiming to extract timing of key stages. Therefore, in the present case of simulated canopy height measurements, we decided to go for a temporal-first approach. For other HTFP techniques such as for example thermographic measurements where spatially correlated measurement errors can be large, a spatial-first approach may be of advantage. It is therefore essential in HTFP to base the modeling decision on the orders of magnitude of the phenotypic variation in relation to the measurement errors.

For the P-spline/QMER method, processing multiple years using a linear mixed model analysis reduced the variance and bias of predictions while slightly increasing the RMSE for *t*PH_stop_ but not for *t*PH_start_. Weighting the first stage further reduced the RMSE. For the second stage, using weights based on estimated variances to approximate the gold standard of a single-stage analysis proved to be of advantage for all traits if using meaningful weights for the first stage as well. These findings indicate that our assumption about dynamic modeling was justified: the spatio-temporal correlation caused by unconsidered covariates yields spatially correlated intermediate traits 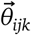. Nevertheless, using a two-stage temporal-first approach with an *AR*(1) ⊗ *AR*(1) autocorrelation structure in the linear mixed model part outperformed the stage-wise approach for *t*PH_start_ and to some extent for *t*PH_stop_.

Overall, the differences between the number of stages (2-stage versus 3-stage) and the order of stages (temporal-first versus spatial-first) were much larger than the differences between the weighting options. Consequently, providing weights may be beneficial but not essential if evaluating real-world experiments with noticeable phenotypic variations.

Nevertheless, using poor error variance estimates to obtain weights may adversely affect the analysis outcome (Cochran, 1954;Rao et al., 1981). A preliminary attempt of us to use posterior distribution simulation based error estimations failed due to a lack of robustness. In contrast, spline predictions have proven to be useful as they allowed a simple and robust estimation of standard errors. We like to emphasize that the recommendation to use spline predictions-based error estimations by assuming proportionality of weights represents a starting point only. Further research is needed to improve the estimation of weights.

Providing robust and reusable analysis routines represented an essential objective of the proposed approach. The resulting generalization requirements may be in conflict with well-established analysis principles. This conflict became well visible when formulating a linear mixed model for Stage 2: The philosophy “analyse-as-randomised” would require to include all randomization factors—e.g., incomplete blocks—in the analysis. A generalized model as used in this work that includes besides a smooth bivariate surface just row and range effects is certainly less efficient, but may nevertheless be suitable to draw correct conclusions on the outcome of the experiment. Proposing a robust and reusable processing workflow therefore always represents a trade-off between generalization and most efficient modeling.

### 4.2. Intermediate trait categories

In this study, we proposed three different trait categories: (1) Timing of key stages, (2) quantities at defined time points or periods, and (3) dose-response curves. A fundamental difference between traits of the first two categories and dose-response curve traits is how they include covariate dependencies. Dose-response curve traits describe an explicit dependency on covariates. In contrast, timing of key stages’ traits include the effects of covariates implicitly through the dependency on the timescale: Favorable conditions in spring may for example accelerate the development of plants and therefore early key stages. Quantities at defined time points or periods’ traits may show a similar behavior, but here the directions are less clear: Early jointing in cereals due to favorable conditions in spring may for example reduce the early canopy cover in the corresponding phase because of a reduced growing time span. Nevertheless, one may also argue that favorable conditions in this reduced time span may increase canopy cover. Both categories have in common that they describe an implicit reaction to a set of covariate courses.

Consequently, to analyze traits of the first two categories, one reduces growing seasons with their characteristic covariate courses to environments (E) and quantifies the influence of genotypes (G) and environments on measured traits in a subsequent G×E analysis (for an overview see van Eeuwijk et al., 2016). In contrast, dose-response curve traits are less affected by—but rather drivers of—G× E. This difference may require differing processing steps. We will cover dose-response curves in a follow-up paper (Roth et al., 2021).

### 4.3. Limitations of dynamic modeling

Clear limitations of the proposed approach became visible: Although all input parameters of the simulation were completely uncorrelated, the extracted traits were to varying extents correlated. The simulation consisted of 500 independent simulation runs, and correlations were aggregated over all runs. Therefore, the observed effects are a systematic result of the extraction methods and should be seen as corresponding limitations. When using P-splines to extract key points of the stem elongation, the estimated start of the stem elongation may be biased by the base temperature of growth. Nevertheless, this effect presumably applies to any other method including the Percentile method, as both early start and low base temperature may result in a comparable phenotype in early stages.

An increased length of the measurement interval may save considerable time and labor costs which may be invested in a larger number of tested genotypes. If aiming to extract timing of key stages, high frequencies are to some extent superfluous if using P-splines, as the spline approach is presumably able to interpolate critical measurement time points. Therefore, one to two measurements a week are sufficient, providing that the total number of measurements does not drop below eight data points (as fitting a shape constrained P-spline using the *scam* package to a time series with less than eight data points becomes challenging in our experience).

### 4.4. Limitations of processing in stages

A salient feature of our suggested approach is to proceed in several stages, starting with an analysis of time series per plot. Because of this feature, our approach does not explicitly account for gross day-dependent errors operating across all plots, although such errors represent an issue in real field data (Kronenberg et al., 2020b). Explicitly accounting for such errors while also modelling the temporal trajectory would require joint spatio-temporal modelling of the time series across all plots simultaneously. There are several approaches for spatio-temporal modelling of environmental data that could be used here. As we are using splines for modelling both the temporal and the spatial dimension, the most immediate option would be to use three-dimensional tensor spline smoothing (Wood, 2017;Verbyla et al., 2018;Pérez et al., 2020). However, most of these are more complex and computationally demanding and as such less suited for a seamless implementation for routine analysis.

## 5. Conclusion

Processing repeated plot-level measurements using a well-defined process and data model revealed insights on best practice in phenomics data handling. The results confirmed that HTFP measurements allow extracting genotype specific timing of key stages and quantities at defined time points. P-splines combined with the QMER method allowed extracting the timing of key stages and quantities at defined time points with a precision that is suitable for plant breeding purposes.

Weighting turned out to be essential if processing HTFP data in stages, and linear mixed model analysis was suitable to account for heterogeneity introduced by covariates not considered. Clear restrictions of the proposed data processing strategy became obvious: Correlations between extracted traits cannot only arise from data, but also from the extraction method itself. Therefore, care has to be taken when interpreting such correlations.

Yet, overall, the scientific community dealing with crop phenotyping has not come up with generally accepted procedures how to organize the workflow from raw data generation to extraction of physiologically meaningful results. This modeling framework is a first step to achieving this aim; not only for the merit of increased scholarly knowledge generation, but in the interest of a more efficient workflow for crop breeding to improve global nutrition aspects in times of climate change.

## Appendix

### 5.1. Table: Simulation input parameters

**Table 2:** Model input parameters for the simulation

|  | Distribution | Values | Sources |
| --- | --- | --- | --- |
| $\theta_i^C$ | $\mathcal{N}(\mu_C, \sigma_C^2)$ | $T_{\min}: \mu_C = 8, \sigma_C = 2$<br>$T_{\text{opt}}: \mu_C = 18, \sigma_C = 2$<br>$r_{\max}: \mu_C = 0.9, \sigma_C = 0.2$ | Kemp and Blacklow (1982)<br>Kemp and Blacklow (1982)<br>Own data |
| $\theta_{jk}^C$ | $\text{AR}(1)_x \otimes \text{AR}(1)_y$ | $\rho_{x\&y} = 0.95, \sigma_{x\&y} = \frac{\sigma_C}{2\sqrt{2}}$ | Velazco et al. (2017) |
| $\theta_i^T$ | $\mathcal{N}(\mu_T, \sigma_T^2)$ | 2016, $t\text{PH}_{\text{start}}: \mu_T = 108, \sigma_T = 2.8$<br>2017, $t\text{PH}_{\text{start}}: \mu_T = 103, \sigma_T = 3.0$<br>2018, $t\text{PH}_{\text{start}}: \mu_T = 101, \sigma_T = 3.1$ | Kronenberg et al. (2020a)<br>Kronenberg et al. (2020a)<br>Own data |
| | | 2016, $t\text{PH}_{\text{stop}}: \mu_T = 165, \sigma_T = 2.5$<br>2017, $t\text{PH}_{\text{stop}}: \mu_T = 162, \sigma_T = 3.5$<br>2018, $t\text{PH}_{\text{stop}}: \mu_T = 158, \sigma_T = 4.0$ | Kronenberg et al. (2020a)<br>Kronenberg et al. (2020a)<br>Own data |
| $\theta_{jk}^T$ | $\text{AR}(1)_x \otimes \text{AR}(1)_y$ | $\rho_{x\&y} = 0.95, \sigma_{x\&y} = \frac{\sigma_T}{2\sqrt{2}}$ | Velazco et al. (2017) |
| $e_{jkt,1}$ | $\text{AR}(1)_t$ | $\rho_m = 0.7, \sigma_m = 0.01$ | Roth et al. (2020) |
| $e_{jk}$ | $\text{AR}(1)_x \otimes \text{AR}(1)_y$ | $\rho_{e,x\&y} = 0.7, \sigma_{e,x\&y} = \frac{\sigma_m}{50}$ | Assumption |
| $e_{jkt,2}$ | $\mathcal{N}(\mu_{e,k}, \sigma_{e,k}^2)$ | $\mu_{e,k} = 0, \sigma_{e,k} = \frac{\sigma_m}{100}$ | Assumption |

### 5.2. Table: Median bias, variance and root-mean squared errors for the P-spline/QMER method

**Table 3:**
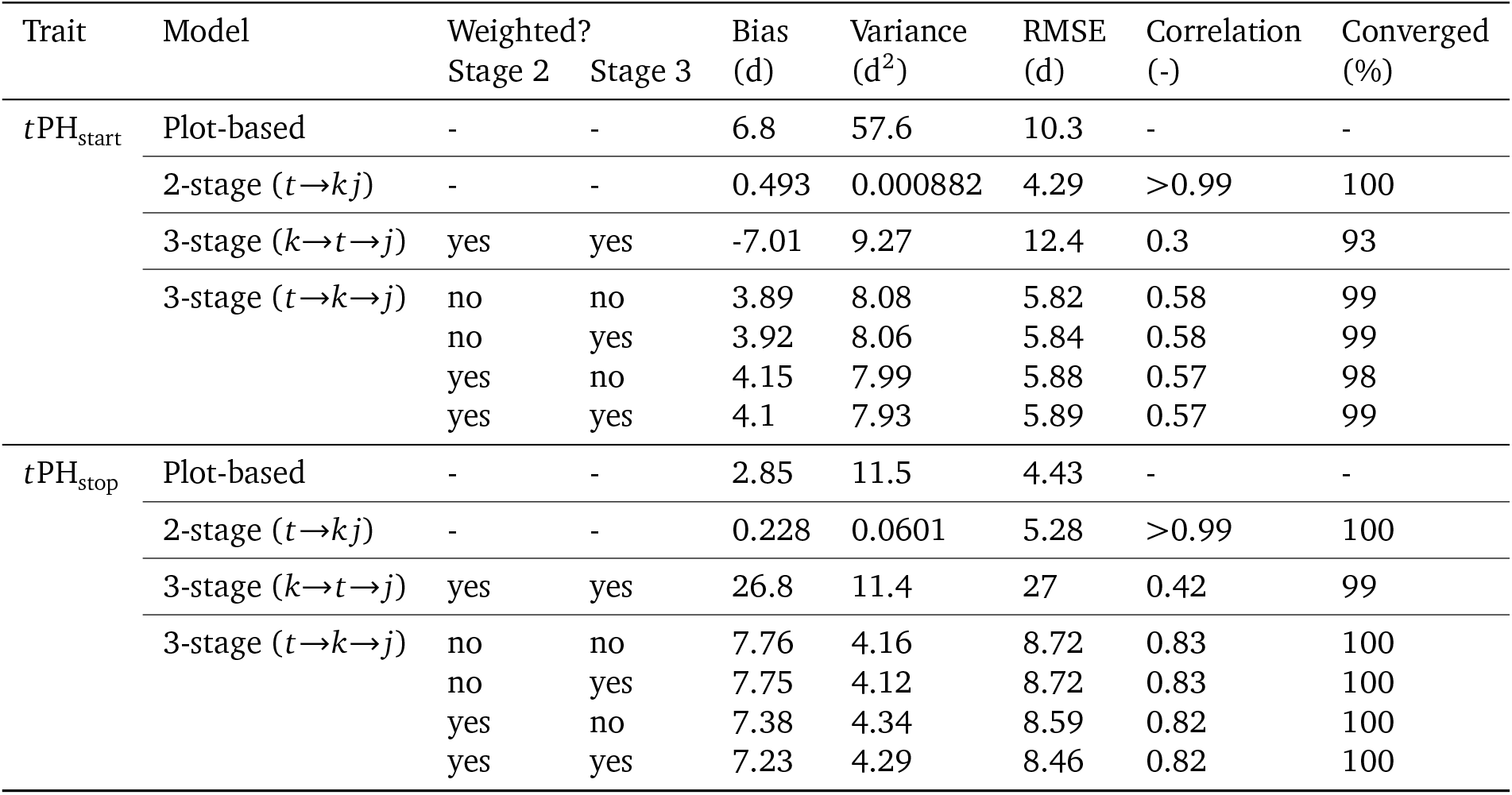
Genotype based bias, variance, root-mean squared error (RMSE), and Pearson’s correlation for the key stages obtained using the P-spline/QMER method, with weighting as option for the second and third stage of the spatial-first (*k*→*t*→*j*) and the temporal-first (*t*→*k*→*j*) three-stage model, and weighting as option for the second stage of the temporal-first (*t*→*kj*) two-stage model. Results report the median values over the 500 simulated datasets. For sake of completeness, plot-based median values for the P-spline/QMER method are reported as well.

### 5.3. A thought on weighting for traits of the second category (timing of key stages)

Splines can be thought of as polynomials, or other functions that are linear in the regression parameters, pieced together at the knots. Thus, to gain some insight, we here consider a quadratic polynomial as a simple concrete example: *f* (*t*) = *μ* + *β*_1_*t* + *β*_2_*t*^2^. We observe data *y_i_*(*t*) = *f* (*t*) + *e_i_* (*i* = 1,…, *n*), where 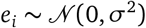. The model is linear and can be written in general for as *y* = *Xβ* + *e*, where 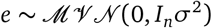. Parameters can be estimated by ordinary least squares using 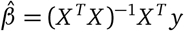 with

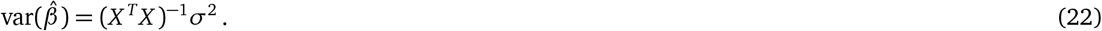

A prediction at a particular value of *t* is obtained from 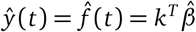 with *K^T^* = (1 *t t*^2^), and this has variance

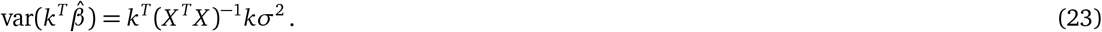

By way of illustration assume that the aim is to find the value of *t* at which the response *f* (*t*) is maximized. For simplicity, we take for granted that a maximum indeed occurs in the relevant range for *t*. At the maximum, the slope of the curve, i.e. the first derivative equals zero. This can be used to determine the optimal input level: 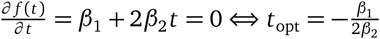. This can be estimated by 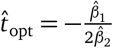.

Now what can be said about the variance of this estimator, which would be needed for weighting? Here, we may use the delta method (Johnson et al., 1993) to find

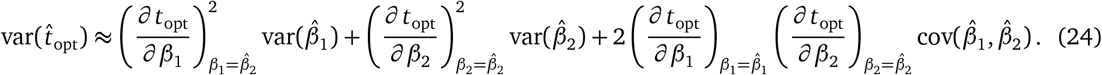

From Equation 22, this is a linear function of *σ*^2^. Now Equation 23 is also linear in *σ*^2^. This suggests that the weights for 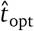 will be positively associated with those for 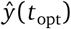. Exact proportionality cannot be expected, however, because whereas *k^T^*(*X^T^X*)^−1^*k* in Equation 23 is constant across plots, the variance in Equation 24 depends on regression parameters that are plot-specific. However, as long as these parameters are not very variable between plots, the association between weights for 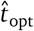 and 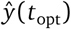 may be expected to be positive and high.

For illustration, we have considered the problem if locating the optimum of a quadratic curve. Note that for different inverse prediction problems, similar expressions would result for the approximate variance, all of which are linear in *σ*^2^. For example, the quadratic is the first derivative of a cubic model, and the optimum of the derivative corresponds to the turning point of the cubic model. Similarly, we can consider the point at which a linear or quadratic model crosses the abscissa, which via the delta method yields an approximate variance of the point estimate that is linear in *σ*^2^.

### 5.4. Averaged parameters of repeated curves versus the curve at the cross-sectional average

To better understand the effect of applying a spatial-first versus a temporal-first approach when extracting timing of key stages, we simplified the simulation used in this study to a logistic growth curve,

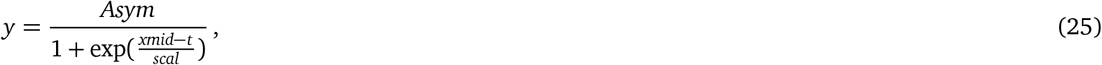

where *Asym* is the asymptote of the curve, *xmid* the inflection point, and *scale* the slope at the inflection point. By varying *xmid* around a mean value 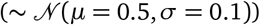 while keeping *Asym* fixed to 1 and *slope* to 0.1, we simulated synchronized but early or delayed *t*PH_start_ and *t*PH_stop_ of phenotypes of the same genotype (Figure 10a, black lines). We then computed two “average genotype” curves, one at the parameter mean (*xmid* = 0.5) (corresponding to a temporal-first approach), and the other one based on cross-sectional averages of individual curves (corresponding to a spatial-first approach) (Figure 10a, blue and red lines). We then extracted *t*PH_stop_ based on the QMER method from the individual curves (Figure 10b, grey boxplot) and from the two average curves (Figure 10b, blue and red lines).

**Figure 10:**
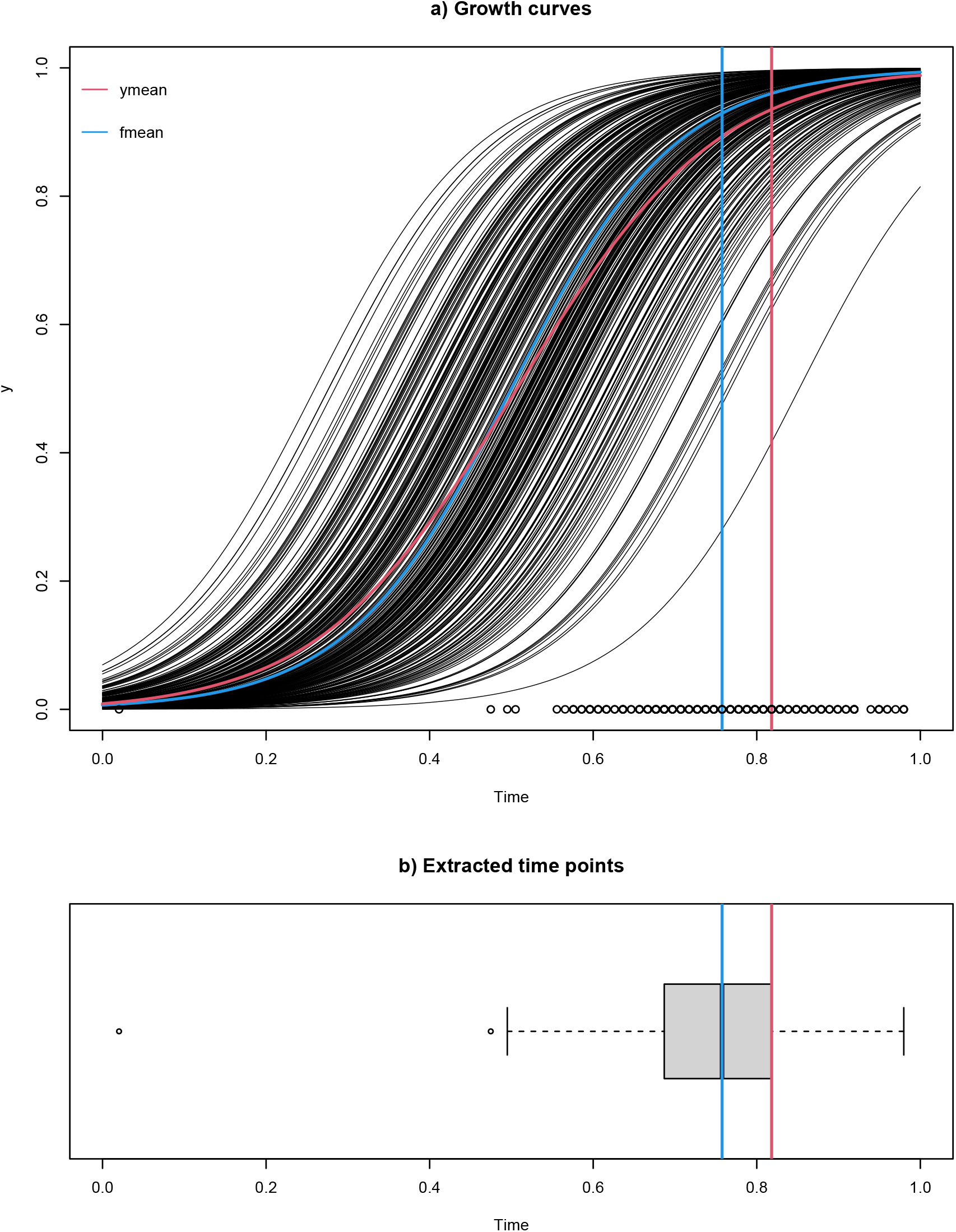
Example of 100 logistic growth curves with fixed asymptote (1.0) and slope (0.1) and Gaussian distribution centered inflection points at 0.5 (a) and the QMER based extracted time points for the end of growth (b), simulating the phenotypic variation of one single genotype. Results of three different extraction approaches are indicated: Applying the QMER method to individual curves (black circles in (a) and grey boxplot in (b)), to a curve that represents the cross-sectional average (red lines in (a) and (b), ymean), and to a curve that is based on averaged curve parameters (asymptote at 1.0, slope at 0.1, inflection point at 0.5) (blue lines in (a) and b), fmean). The ymean approach is comparable with a spatial→temporal approach, the fmean approach with a temporal→spatial approach.

As expected, *t*PH_stop_ extracted at the parameter mean curve was around the average of the *t*PH_stop_’s of the individual curves. However, for the cross-sectional curve, the extracted *t*PH_stop_ was far from this average.

We therefore see indications that when applying the QMER method in a temporal-first or spatial-first approach, the same method will, depending on the order of spatial and temporal modelling, estimating different things. Furthermore, we see indications that the temporal-first approach is closer to the data generating mechanism. Ultimately, the question arises which of the average curves one thinks is a better descriptor of what one would call the “genotypic” curve. A researcher with a biological background may argue (as we did in this manuscript) that when sampling a number of phenotypes of the same genotype on a field, the chance to sample a phenotype with a growth curve close to the blue one in Figure 10a is highest, and therefore consider this as the “average” genotype. Nevertheless, a more statistical view is that, on average, the observed phenotypes have a growth curve relating to the red one in Figure 10a. Interestingly, such questions would most probably also arise when specifying a full spatio-temporal model.

## Acknowledgement

We acknowledge Helge Aasen, Lukas Kronenberg and Norbert Kirchgessner (ETH Zurich) for feedback on an early version of the manuscript. Furthermore, we thank the Informatik Support Gruppe (ISG) D-HEST of ETH Zurich for spontaneously helping us out with short-term computing capacity required to complete the simulation runs. Finally, we like to thank the two anonymous reviewers for their exceptionally detailed and thoughtful feedback that helped to significantly improve the manuscript.

## CRediT authorship contribution statement

**Lukas Roth**: Conceptualization, Methodology, Software, Formal analysis, Visualization, Writing - Original Draft. **María Xosé Rodríguez-Álvarez**: Software, Writing - Review & Editing **Fred van Eeuwijk**: Writing - Review & Editing. **Hans-Peter Piepho**: Conceptualization, Methodology, Writing - Original Draft, Review & Editing. **Andreas Hund**: Conceptualization, Supervision, Project administration, Funding acquisition, Writing - Review & Editing.

## Data availability

Data and source code that support the findings of this study are openly available in the ETH gitlab repository at https://gitlab.ethz.ch/crop_phenotyping/htfp_data_processing and archived in the ETH research collection (http://doi.org/10.5905/ethz-1007-385).

